# Smad4 signaling establishes the somatosensory map of basal vomeronasal sensory neurons

**DOI:** 10.1101/738393

**Authors:** Ankana S. Naik, Jennifer M. Lin, Ed Zandro M. Taroc, Raghu R. Katreddi, Jesus A. Frias, Morgan Sammons, Paolo E. Forni

## Abstract

The accessory olfactory system is a unique model that can give insights on how the neurons can establish and maintain their identity, and connectivity. The vomeronasal organ (VNO) contains two distinct populations of vomeronasal sensory neurons (VSNs) each with specific innervation patterns to the accessory olfactory bulb (AOB). Though morphogenic signals are critical in defining various neuronal populations, the morphogenic signaling profiles that influence each VSN population remains unknown. Here, we found a pronounced BMP signaling gradient within the basal VSNs. By generating Smad4 conditional mutants, we disrupted canonical TGF-β/BMP signaling in maturing basal VSNs and in all mature VSNs. We show that Smad4 loss-of-function in immature basal neurons leads to a progressive loss of basal VSNs, reduced activation of the remnant basal VSNs, and aberrant glomeruli formation in posterior AOB. However, Smad4 ablation in all mature VSNs does not affect neuronal activity nor survival but causes aberrant glomeruli formation only in the posterior AOB. Our study reveals that Smad4 signaling plays a critical role in mediating development, function, and circuit formation of basal VSNs.

## Introduction

Neurons form complex and conserved neuronal circuits by innervating the dendritic compartments of specific target neurons. One current goal in neuroscience is to delineate the underlying mechanisms that establish and maintain neuronal diversity, identity, and connectivity. The vomeronasal organ (VNO) is a specialized vertebrate olfactory subsystem used to detect pheromones (Cloutier et al., 2002; Dulac, 2000; Isogai et al., 2011; Mombaerts et al., 1996). As the main olfactory epithelium, the vomeronasal neuroepithelium continually generates new neurons (de la Rosa-Prieto et al., 2010). As newly formed neurons mature, they innervate dendrites of specific second-order neurons in the accessory olfactory bulb (AOB) (Mombaerts et al., 1996). The sensory epithelium of VNO in mice is composed of vomeronasal sensory neurons (VSNs) that selectively express receptors encoded by one of the two vomeronasal receptor (VR) gene families: V1r and V2r. These non-overlapping V1r- and V2r-expressing populations of VSNs bind different ligands, project to different areas of the AOB, and trigger distinct innate behaviors (Chamero et al., 2012). V2r-expressing neurons localize to the basal portion of the vomeronasal epithelium and target posterior regions of the AOB, while V1r-expressing neurons reside in the apical portion and project to anterior regions of the AOB.

Olfactory sensory neurons (OSNs) in the main olfactory system that express the same olfactory receptor gene target the same glomeruli in the olfactory bulb (OB). However, in the vomeronasal system, VSNs expressing the same receptor coalesce onto multiple glomeruli within spatially conserved regions of the AOB (Belluscio et al., 1999; Del Punta et al., 2002). Recent evidence indicates that unidentified spatial cues in the nasal area likely influence OSN gene choice and target specificity (Coleman et al., 2019). Our current knowledge of the underlying signals that establish the spatial (apical-basal) identity of VSNs and delineates synaptic partners remains limited.

Molecules belonging to the Transforming Growth Factors (TGF) super-family, such as TGFβ, Activin, and bone morphogenic proteins (BMPs), control nervous system growth, differentiation, axonal growth patterning, and network formation (Banerjee and Riordan, 2018; Le Dreau et al., 2012; Lee et al., 2000; Liem et al., 2000; Liem et al., 1995; Schmidt et al., 1995). Retrograde BMP signaling controls neuromuscular junction synaptic growth and cytoarchitecture in drosophila (Ball et al., 2010; Piccioli and Littleton, 2014). BMP signaling also contributes to establishing neuronal identity and defining somatosensory map formation in the rodent trigeminal nerve (Ball et al., 2010; Banerjee and Riordan, 2018; Berke et al., 2013; Fuentes-Medel and Budnik, 2010; Hegarty et al., 2013; Hodge et al., 2007; Liao et al., 2018; Piccioli and Littleton, 2014).

Whether molecules of the TGF-β family play roles in defining specific neuronal identities and synaptic circuit formation of the VNO with the brain has not been explored thus far. Smad4 is a central signaling molecule in canonical TGF-β/BMP intracellular signaling. Upon ligand-receptor binding, Smad4 forms a complex with phosphorylated R-Smads (Smad1, 5, 8 for BMPs or with Smad2, 3 for TGF-β) that translocate to the nucleus to activate gene transcription (Shi and Massague, 2003). BMP ligand’s affinity to extracellular matrix components such as collagen IV (Col-IV) define spatial BMP signaling gradients (Bunt et al., 2010; Garamszegi et al., 2010; Paralkar et al., 1991; Paralkar et al., 1992; Wang et al., 2008).

Here, we characterized the role of Smad4 mediated signaling to define neuronal identity, circuit formation, and/or homeostasis of VSNs using the AP-2εCre mouse line to conditionally ablate Smad4 expression in immature basal VSNs and OMPCre mice to ablate Smad4 in mature apical and basal VSNs. We show that loss of Smad4 dependent intracellular signaling compromises functionality, survival, and proper circuit formation of basal VSNs to the pAOB.

## Results

### Localization of active BMP signaling in basal VSNs

We analyzed mRNA-seq data from postnatal VNO and discovered a source of multiple molecules of the TGF-β family (GSE134492). We validated gene expression of select genes via RT-PCR (Fig.1 A,B). By analyzing the expression of BMPs in the VNO using in situ hybridization against BMP4 and BMP6, we revealed BMP expression both in apical and basal territories of the VNE (Fig. 1C,D). Extracellular matrix components such as Collagen IV (Col IV) may participate in establishing active BMP signaling by sequestering or immobilizing morphogens and facilitating receptor binding (Bunt et al., 2010; Garamszegi et al., 2010; Paralkar et al., 1991; Paralkar et al., 1992; Wang et al., 2008). Col IV and PECAM immunostaining highlighted Collagen IV expression in the basement membrane of the VNO and around the PECAM+ vasculature that invade the basal positions of the VNO (Fig.1 F1-2).

**Fig. 1.**
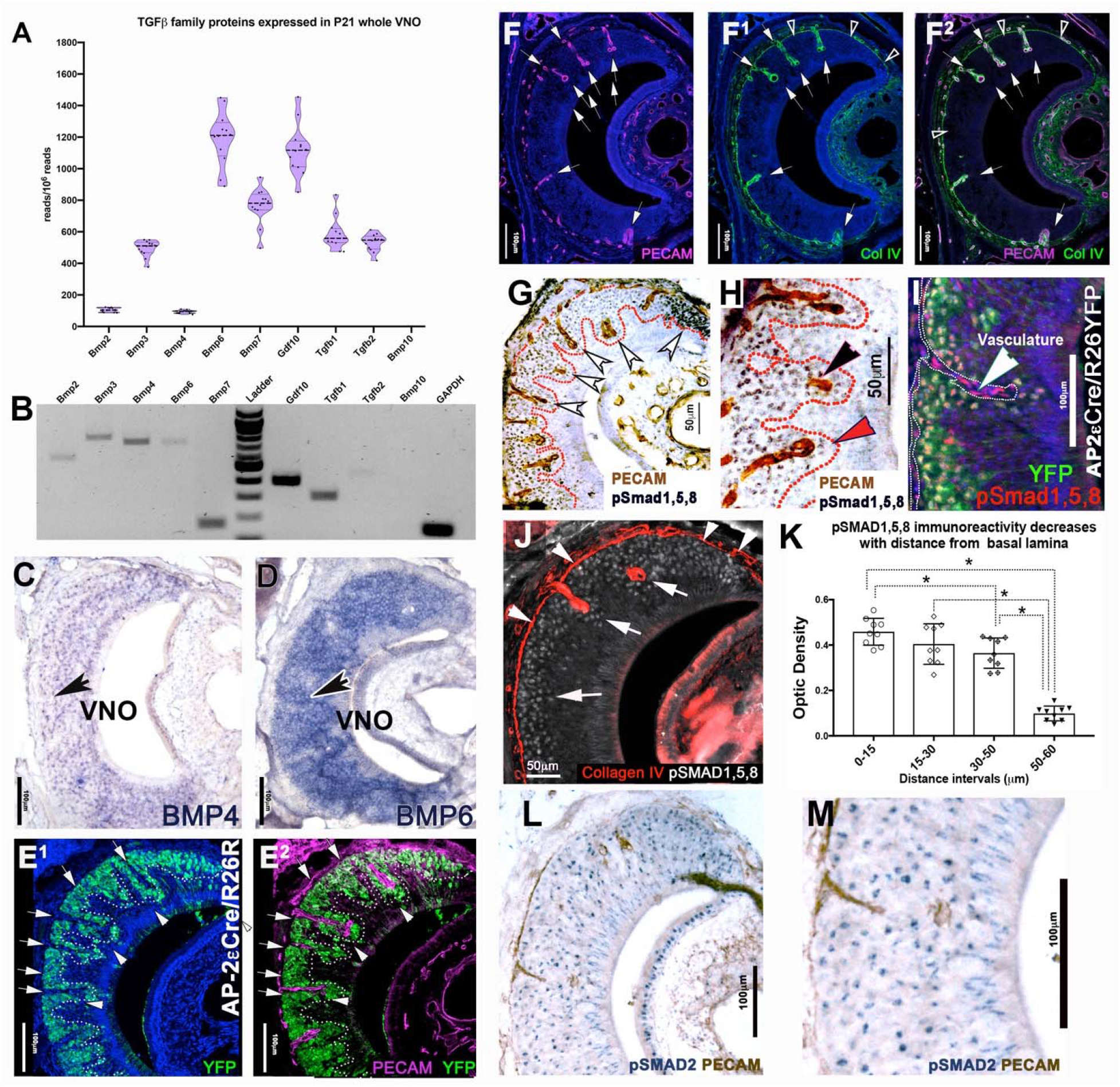
TGF-β/BMP signaling in the Vomeronasal organ. A) Transcript abundance of select BMP and TGFβ molecules according to RNASeq analysis. B) RT-PCR confirmation of select BMP and TGFβ molecules from RNASeq results. C) In situ hybridization for BMP4 and BMP6 (D) on P15 VNO shows uniform distribution of ligand in the neuroepithelium. E1-2) Immunofluorescence against AP-2εR26YFP (P15) lineage tracing (green), the vasculature marker PECAM (magenta) and DAPI (blue) shows a close spatial association of the YFP+ basal VSNs to vasculature in the VNE (white arrowheads). F1-3) Immunofluorescence on WT (P21) against PECAM (Magenta) and Collagen IV (ColIV, green) with a DAPI (blue) counterstain shows the basal lamina positive for Col IV (black arrowheads) encapsulating the invading vasculature (white arrows) G) Immunohistochemistry on WT (P15) shows p-Smad1,5,8 immunoreactivity (blackish-gray) in VSNs proximal to the PECAM positive (brown) vasculature transducing BMP signaling. H) Magnification of (G), black arrow pointing the PECAM positive vasculature and red arrow showing the p-Smad1,5,8 positive VSNs transducing BMP. I) p-Smad1,5,8 (Red) and AP-2εCre/R26RYFP lineage tracing (green). All the basal VSNs, positive for AP-2ε-driven Cre recombination (YFP) have strong active BMP signaling, vasculature positive for only p-Smads1,5,8 (white arrow). J) Collagen IV immunostaining (Red) highlights the basement membrane (white arrows) and p-Smad1,5,8 cells (gray) have stronger immunoreactivity proximal to the sources of collagen IV (white notched arrows). K) p-Smad1,5,8 optic density (OD) after DAB staining at 4 distance intervals from the basement membrane in the VNO, unpaired t-Test p (*<0.05), +/-SEM, (n=3 animals; individual points represent the average of 30 cells counted per interval/sections; 3 sections/animal). L) Immunostaining anti p-Smad2 (Blue) shows TGFβ signaling in the VNO with no apparent gradient with respect to PECAM (brown) positive vasculature M) Magnification of (L) showing uniform expression of p-Smad2 in the VNE.

We then performed immunostaining against p-Smad1,5,8 and p-Smad2 to monitor the activity of BMP and TGF-β signaling pathways in the VNE. P-Smad1,5,8 immunostaining showed immunoreactivity in PECAM positive endothelial cells (Fig.1 I) and in VSNs in the basal territories proximal to PECAM+/Col IV+ vessels and basal lamina (Fig.1 G,H). Using sections from AP-2εCre^+/-^/R26R lineage traced animals (Lin et al., 2018), where basal cells are genetically traced after AP-2ε expression (Fig. 1 I), double immunofluorescent staining against p-Smad1,5,8 showed that nearly all basal VSNs were positive for active BMP intracellular signaling beginning at the marginal zone (Fig. 1 I, S1 B,D). We detected only background levels of p-Smad1,5,8 immunoreactivity in the apical territories. In contrast, p-Smad2 immunostaining indicated active TGF-β signaling in both apical and basal territories (Fig.1 L,M).

Since the pattern of BMP signaling appeared stronger in cells in the basal territories proximal to Col IV rich basement membrane and vasculature (Fig. 1 G,H), we performed a densitometric analysis for p-Smad1,5,8 cell immunoreactivity in relation to distance from the basement membrane. We found a negative correlation in the expression of p-Smad1/5/8 and the distance from the basement membrane (Fig.1 J K). Based on these data, we propose that basal VSNs actively transduce BMP signaling and that there is gradient of active BMP intracellular signaling within the basal region of VNO. We then analyzed the VNE from Arx-1 null mice, which lack an olfactory bulb and have no connection between VSNs and the brain (Taroc et al., 2017). We found a similar pattern of p-Smad1,5,8 immunoreactivity as in control animals (data not shown). These data suggest that the identified p-Smad 1,5,8 basal to apical gradient in the VNO reflects locally produced morphogenic signals rather than retrograde signaling as shown for other systems (Ji and Jaffrey, 2012).

### Maturing vomeronasal sensory neurons show active BMP signaling in marginal regions of the VNO

In addition to the basal-apical BMP signaling gradient, our densitometric analysis also revealed significantly stronger p-Smad1,5,8 immunoreactivity in basal marginal regions of the VNO compared to medial regions (Fig. S1 B). The marginal regions contain immature basal (GAP43+; AP-2ε+; OMP-) and immature apical VSNs (GAP43+; Meis2+; OMP-) (Fig. S1,C,D,F,G, Use Fig. S1A as a reference) (Brann and Firestein, 2010; Giacobini et al., 2000). We then quantified pSmad1,5,8, AP-2ε and GAP43 immunoreactivity (Fig. S1 B-D). We observed a higher level of expression for all three markers in the marginal regions of the VNO compared to medial portions where neurons reach maturity (OMP+) (Fig. S1 B,C,F). In contrast to the pattern of active BMP signaling, immunostaining and quantification for active TGF-β signaling p-Smad2 and apical marker Meis2 did not differentiate marginal and medial regions of the VNE (Fig. S1 E,G). Based on the spatial correlation between high levels of active BMP signaling (p-Smad 1,5,8) and the basal VSN specific transcription factor (AP-2ε) in immature VSNs (GAP43) (Fig. S1 B,C,D), we speculated BMP may contribute to the maturation process of basal VSNs.

### Conditional ablation of Smad4 disrupts TGF-**β**/BMP signaling in basal VSNs

To investigate the role of canonical TGF-β/BMP signaling in differentiating basal VSNs, we exploited AP-2εCre mice as a genetic entry point to conditionally ablate Smad4^flox/flox^ (Yang et al., 2002) in immature (GAP43+) basal VSNs (Lin et al., 2018). We validated AP-2εCre mediated ablation at postnatal age P15. The VNO at this stage is functional and close to its final size, but still retains a considerable amount of immature neurons generated during the postnatal proliferation peak (Wakabayashi and Ichikawa, 2007). We validated the ablation of functional Smad4 with two approaches. We performed in situ hybridization using an RNA probe against exon 8 of the Smad4 gene, which is flanked by LoxP sites (Yang et al., 2002), and immunohistochemistry against Smad4 (Benazet et al., 2012; Yang et al., 2002). Smad4 mRNA and protein expression analysis on Smad4^flox/flox^ controls, triple heterozygous mutants AP-2εCre^+/-^/Smad4^WT/flox^/R26^YFP+/-^ and traced conditional KOs AP-2εCre^+/-^/Smad4^flox/flox^/R26^YFP+/-^ verified Smad4 ablation in the cells that underwent Cre mediated recombination (Fig 2 D-F2).

**Fig. 2.**
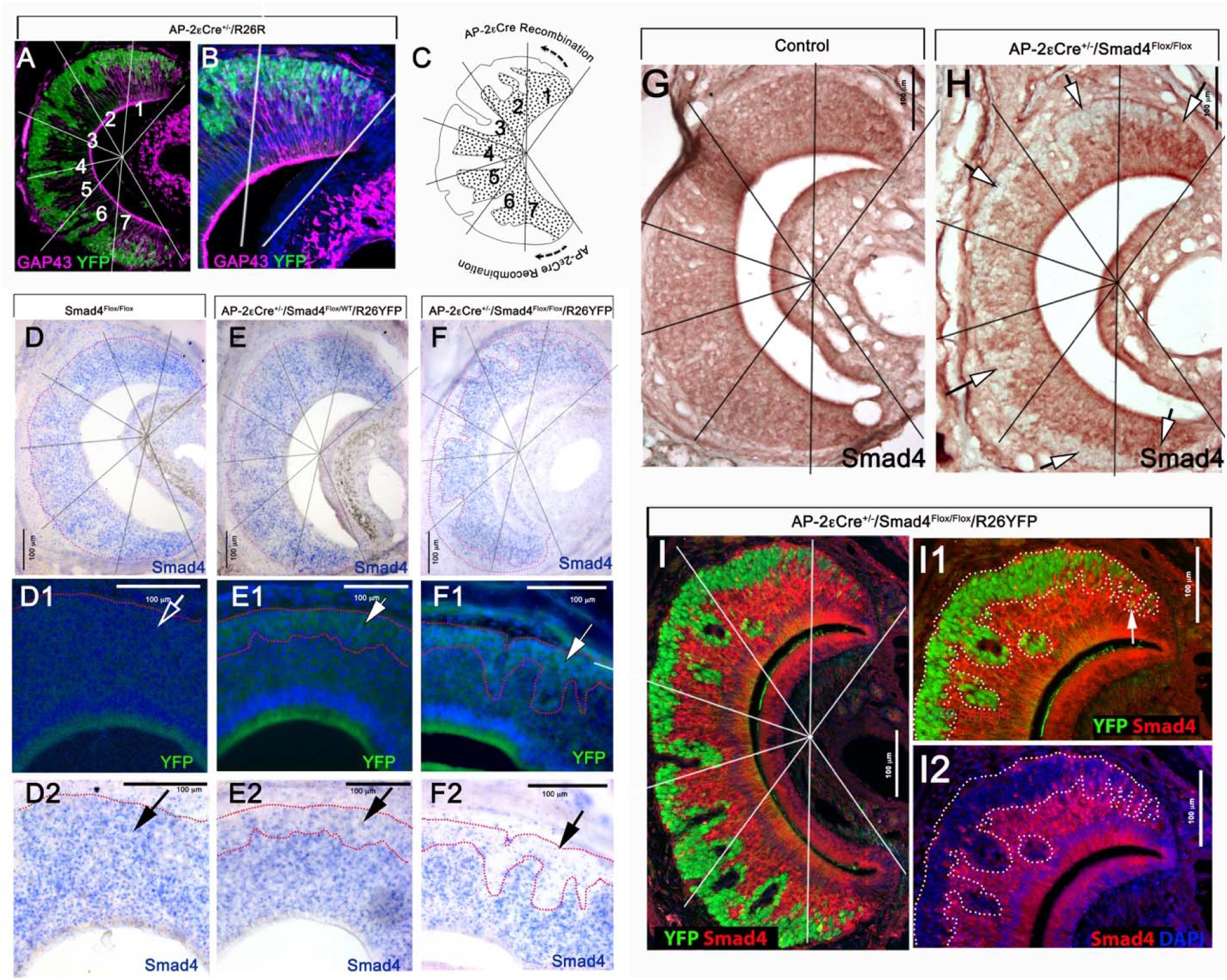
Smad4 ablation in differentiated immature basal VSNs-. A,B) Double immunostaining on AP-2εCre^+/-^/R26YFP (P15) for the immature VSN marker GAP43 (magenta), YFP (green), and DAPI counterstain (blue) highlighting AP-2ε driven Cre recombination in basal VSNs. The recombination occurs specifically in immature basal VSNs. B) Magnification of A. C) Cartoon illustrating AP-2ε driven Cre recombination only in basal VSNs. D,E,F) In situ hybridization for exon 8 of smad4 in P15 Control (D) P15 Smad4 heterozygous tracing control(E) and in P15 Smad4 homozygous traced cKO (F). D1,E1,F1) AP-2ε driven recombination marked by YFP immunostaining in P15 Control (D1) P15 Smad4 heterozygous tracing control(E1) and in P15 Smad4 homozygous traced cKO (F1). D2, E2,F2) YFP immunostaining highlighting AP-2ε driven recombination and Smad4 exon8 ISH showing uniform expression of Smad4 exon8 transcript in the VNE in P15 Control (D2) Lower expression of Smad4 exon 8 transcript in lineage traced basal VSNs in comparison to apical VSNs in P15 Smad4 heterozygous tracing control (E2) and almost no expression of Smad4 exon 8 transcript in traced basal VSNs in P15 Smad4 homozygous traced cKO (F2). G,H) Immunostaining for Smad4 in Control(G) and cKO (H). White arrows indicate complete ablation of Smad4 in basal VSN. I,I1,I2) YFP immunostaining (green) highlighting AP-2ε driven recombination and Smad4(red) immunostaining in P15 Smad4 homozygous traced cKO.

In Smad4^flox/flox^ control animals, visual observation indicated comparable Smad4 mRNA levels in both basal and apical territories of the VNO (Fig.2 D-D2). In triple heterozygous AP-2εCre^+/-^/Smad4^flox/WT^/R26^YFP+/-^ mice, we detected weaker Smad4 expression in basal domains compared to apical regions, likely reflecting the loss of one allele (Fig. 2E-E2). In the conditional KO, the majority of cells positive for AP-2εCre recombination in the basal domain marked by YFP were negative for Smad4 transcript expression (Fig.2 F-F2). Using immunolabeling against Smad4, we confirmed the loss of functional Smad4 protein expression in basal territories starting at the marginal regions (Fig. 2 G,H). Using Smad4 and YFP double-immunolabeling on sections from AP-2εCre^+/-^ /Smad4^flox/flox^/R26^YFP+/-^ mice, we quantified the recombination efficiency of approximately 98% ± 0.716% (Fig. 2 I-I2) starting at the marginal zones of the VNO, which contains immature neurons (Use Fig.S1 A,B as a reference).

### AP-2**ε**Cre driven Smad4 ablation shows no clear abnormalities at 2 weeks after birth

To test the effects of Smad4 conditional ablation in maturing basal VSNs, we analyzed the cell composition and maturation state of cells in the VNE at P15. At this stage, we found a small but significant increase in the number of proliferative (Ki67+) cells, which suggests a compensatory response to epithelial cell loss (Lin et al., 2018), (Fig. 3 A,B,C). However, we did not detect significant changes in the number of cells positive for the apoptotic marker cleaved caspase3 (Fig. 3 D,E,F). Quantification of immunostaining against Gαo indicated that the localization and number of basal VSNs within the VNO remained unchanged in AP-2εCre^+/-^/Smad4^flox/flox^ (AP-2εCre/Smad4cKO) mice. At P15, we also found no changes in the expression of the basal receptor V2R2 (Fig. 3 G-L). However, we did observe slightly more cells immunoreactive for Growth Associated Protein 43 (GAP43) throughout the VNE of the AP-2εCre^+/-^/Smad4^flox/flox^ mice (Fig 3 M-P). These data indicate that Smad4 ablation in maturing basal VSNs as assayed at P15 does not significantly compromise the expression of basal specific proteins and neuronal position of VSNs in the epithelium or their survival.

**Fig. 3:**
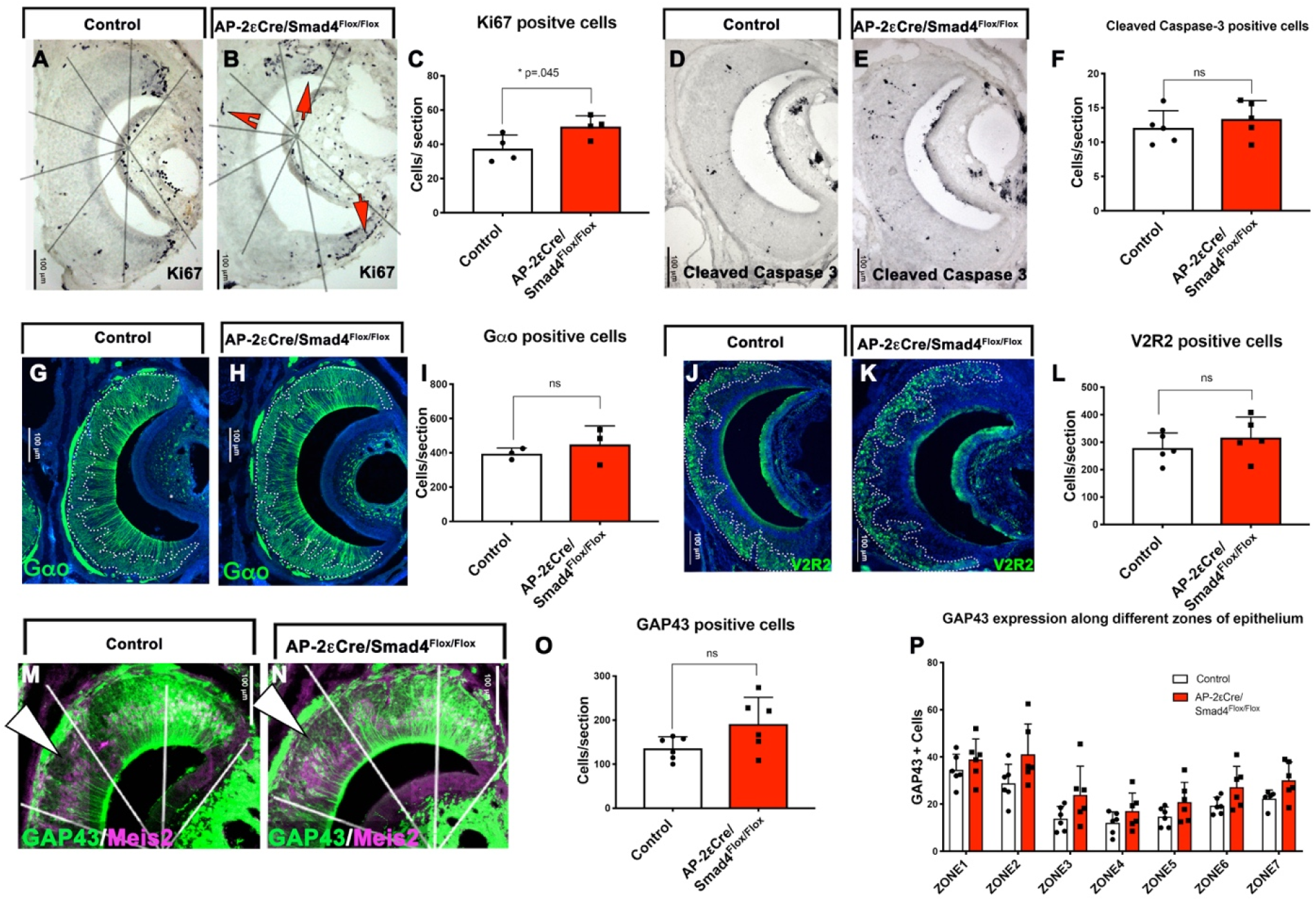
Characterization of AP-2εCreSmad4 at P15 revealed minute changes in VSN development: A,B) Immunostaining for Ki67 positive proliferative cells in control (A) and cKO (B). C) Significant increase in proliferative cells was observed in the cKO(n=4). D,E) Immunostaining for cleaved caspase 3 in control(D) and cKO(E). F) No significant difference was observed in rate of apoptosis between the control and cKO(n=5). G,H) Immunostaining for basal marker Gαo in control (G) and cKO (H). I) No significant difference was observed in number of Gαo positive cells between control and cKO(n=3). J,K) Immunostaining for V2R2 in control (J) and cKO (K). L) No significant difference for V2R2 positive cells was observed between control and cKO (n=5). M,N) Immunostaining for immature marker GAP43 (green) and apical marker Meis2 (magenta) in control (M) and cKO (N). O) Not statistically significant but an overall increase was observed in number of GAP43 positive immature neurons in cKO(n=6). P) Number of GAP43 positive cells in marginal, intermediate and central zones in control and cKO. Statistical analysis by two tailed unpaired, t-test.

### Conditional Smad4 ablation in maturing basal VSNs produces a severe loss of basal neurons in adult mice

The first two post-natal weeks are critical for the development of the vomeronasal organ and formation of functional synaptic connections to the AOB (Hovis et al., 2012; Roos et al., 1988; Weiler et al., 1999). So, we examined if Smad4 ablation affects the homeostasis and/or connectivity of fully mature basal VSNs. We analyzed AP-2εCre^+/-^/Smad4^flox/flox^ conditional KO (AP-2εCre/Smad4cKO) and relative control (Smad4^flox/flox^) at P60, which is the stage when most VSNs reach maturity and have formed functional connections (Hovis et al., 2012). Analysis of AP-2εCre^+/-^/Smad4cKO animals at P60 revealed a 30% to 50% reduction in the number of VSNs expressing the basal markers Gαo, V2R2, and AP-2ε; however, we did not detect any changes in the number of apical (Gαi2 positive) cells (Fig. 4 A-J). The cell density of basal VSNs significantly increased in Smad4cKO mice compared to controls (+54.72%; p= 0.0017), while no difference in cell density occurred in apical VSNs. We then sought to determine if basal cells in Smad4cKO mice transdifferentiated into apical VSNs (Lin et al., 2018) by performing AP-2εCre^+/-^/R26R lineage tracing on Smad4 conditional mutants (AP-2εCre^+/-^ /Smad4^flox/flox^/R26^YFP+/-^). In contrast to the AP-2ε KOs (Lin et al., 2018) these mutants did not show the presence of VSNs that have transdifferentiated from basal to apical VSNs (Fig. 4 K-L1).

**Fig. 4.**
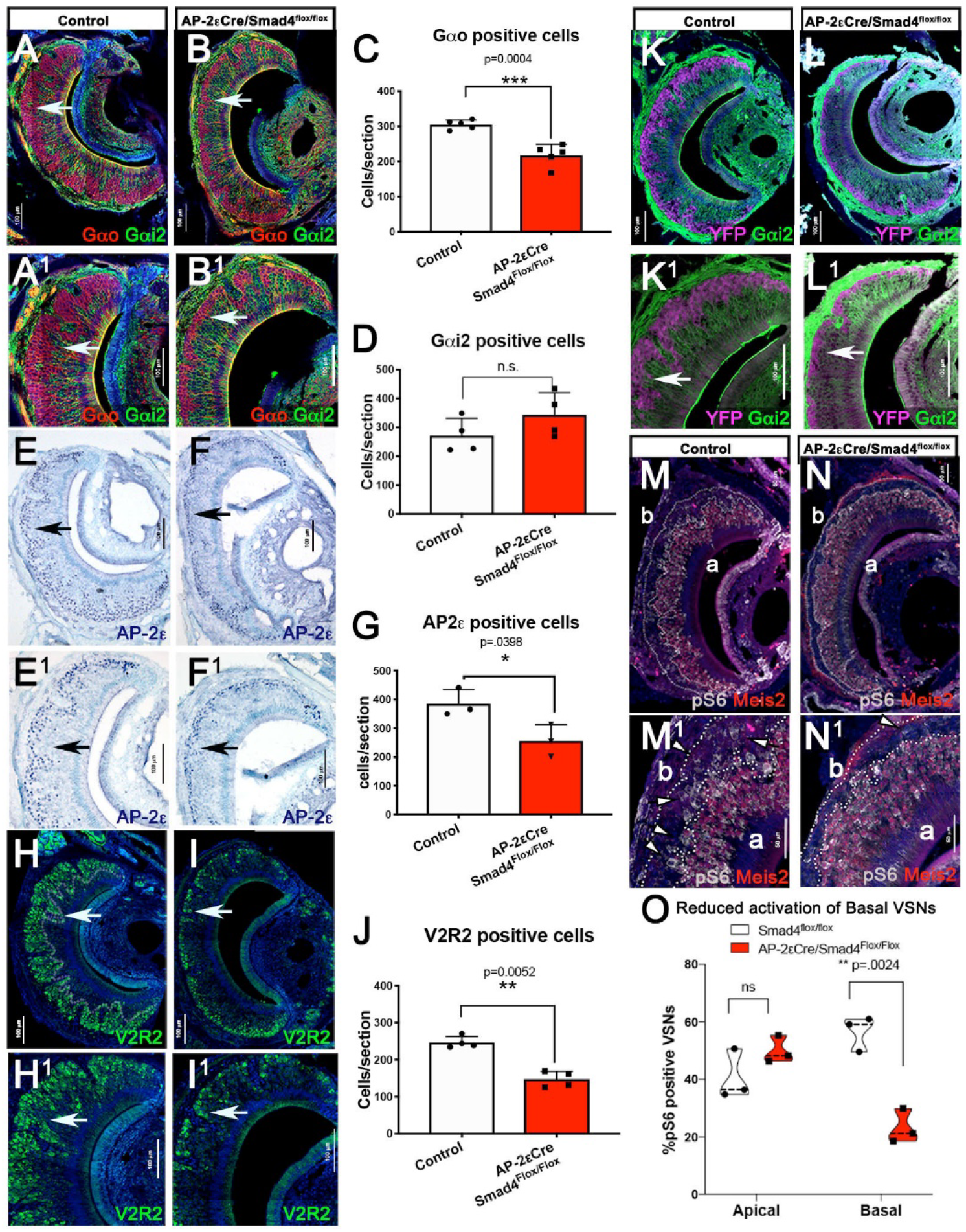
Characterization of AP-2εCreSmad4 at P60 reveals a progressive selection against basal VSNs: A, B) Immunostaining for basal marker Gαo (red) and apical marker Gαi2 (green) on control (A) and cKO (B). A1,B1) magnification of A and B respectively. C) Graph representing significant reduction in number of Gαo positive cells in cKO(n=5). D) Graph representing not a significant increase in Gαi2 positive cells(n=4). E,F) Immunostaining for AP-2 ε on Control (E) and cKO (F). E1,F1) Magnification of E and F respectively. G) Graph representing significant reduction in AP-2ε positive cells in the cKO (n=3). H,I) Immunostaining for V2R2 on Control (H) and cKO (I). H1,I1) Magnification of H and I respectively. J) Graph representing significant reduction in V2R2 positive cells in the cKO (n=4). K,l)Immunostaining for AP-2ε driven recombination (YFP, magenta) and apical marker Gαi2(green) on control (K) and cKO (L). K1, L1) Magnification of K and L respectively. E). M) N) Immunostaining for pS6 (grey) and apical VSN marker meis2 (red) in control (M, magnification M1,M2) and cKO(N, magnification N1,N2).O) Quantification of percentage of pS6 positive apical and basal VSNs in control and cKO show significant reduction in percentage of pS6 positive basal VSNs in cKO (n=3). Statistical analysis by two tailed unpaired, t-test.

By analyzing GAP43 expression at P60 we detected a significant increase in immunoreactivity in the central and the basal region of the VNO of the AP-2εCre^+/-^ /Smad4^flox/flox^ (Fig.S2. A,B,E,F,I,J). By performing Smad4 and GAP43 immunostaining on AP-2εCre^+/-^/R26RYFP control and AP-2εCre^+/-^/R26R^YFP+/-^/Smad4^flox/flox^ mice, we verified that elevated GAP43 immunoreactivity was limited to basal neurons that underwent AP-2εCre mediated recombination (Fig.S2. C-H). Double OMP/GAP43 immunostaining showed GAP43 immunoreactivity in mature basal VSNs in AP-2εCre^+/-^ /Smad4^flox/flox^ mice, while controls only showed sparse double positive cells (data not shown). These data suggest that Smad4 ablation in maturing basal VSNs can still form mature basal VSNs but can compromise aspects of maturation and long-term survival of basal VSNs.

### Smad4 ablation in basal VSNs compromises basal VSNs activation after odor exposure

Exposure to whole male urine activates both basal and apical VSNs (Silvotti et al., 2018). Pheromonal stimulation of VSNs induces sustained phosphorylation of S6 ribosomal protein (pS6), which can serve as an indirect marker of neuronal activity (Silvotti et al., 2018). We assayed VNO function in our Smad4 cKOs by exposing male AP-2εCre^+/-^/Smad4^flox/flox^ and Smad4^flox/flox^ controls to whole male urine and collected the animals 90min after exposure. Analysis of pS6 positive cells showed a significant reduction in activating basal sensory neurons. These data suggest that lack of Smad4 dependent signaling compromises the function of basal VSNs (Fig.4 M-O).

### Smad4 ablation in basal VSNs compromises the formation of glomeruli in the posterior AOB

Our results from AP-2εCre^+/-^/Smad4^flox/flox^ experiments revealed increased GAP43 immunoreactivity in basal VSNs (Fig. S2 A-H). Typically, GAP43 expression occurs in immature neurons that form an axon, undergo synaptic targeting or retain some plasticity (Enomoto et al., 2011; Strittmatter et al., 1990; Strittmatter et al., 1992). To understand if the lack of Smad4 signaling alters axonal targeting, we analyzed VSN projections to the AOB. Immunostaining against the basal axonal marker Robo2 (Prince et al., 2009) and apical axonal marker Nrp2 indicated a ∼50% reduction in the area occupied by Robo2+ (basal neurons) axonal projections to the pAOB of AP-2εCre^+/-^ /Smad4^flox/flox^, consistent with reduced numbers of basal VSNs (Fig. 5E). We did not observe differences in the aAOB (Fig.5E).

**Fig 5.**
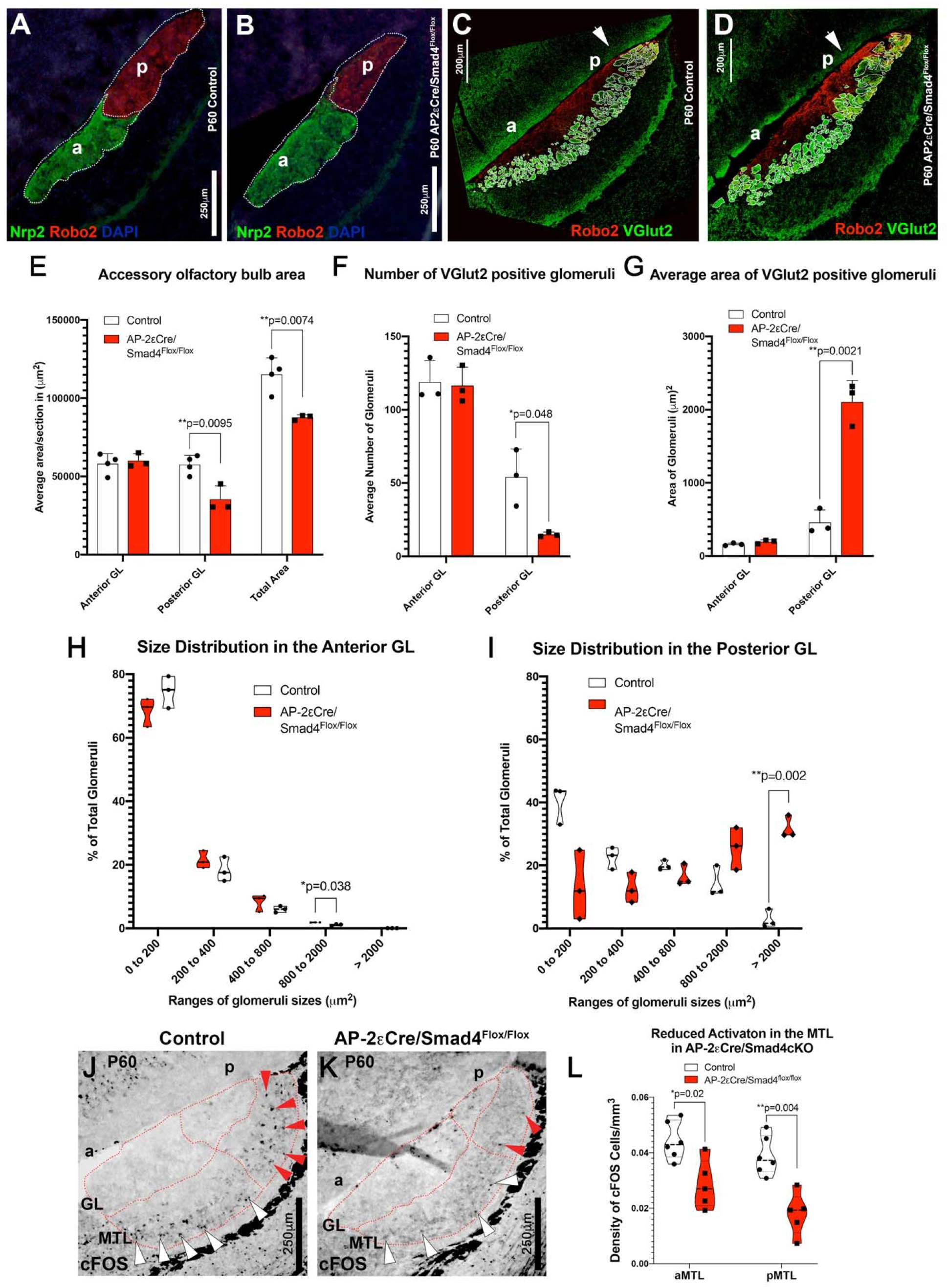
The posterior AOB of AP-2εCre/Smad4cKO is significantly smaller than controls and forms aberrant glomeruli. A,B) Double immunofluorescence against Nrp2 and Robo2 on A) P60 Control and B) P60 AP-2εCre Smad4 cKO. E) Area quantification shows reduced area for the pAOB and reduction in the total area occupied by VSNs’ fibers in the AOB of cKOs but no change in the aAOB(control n=4,cko n=3). C,D) Double immunostaining against Robo2 and VGlut2 highlights the double positive glomeruli in the pAOB of controls (C) and cKOs (D). In the Smad4 cKOs few very large glomeruli were detected in the pAOB, while size and number of the glomeruli in the aAOB appeared to be unaffected. F,G) Quantification of average number and areas of the glomeruli in aAOB and pAOB of controls and AP-2εCre^+/-^/Smad4cKO (n=3). H,I) The percentage of number of glomeruli were binned in area ranges and quantified. The graph shows significant increase in number of glomeruli with increased area in pAOB (n=3). (I) with minor change in aAOB (H). J,K,L) Immunostaining and quantification (L) for cFOS on P60 control (Smad4^flox/flox^) and P60 AP-2εCre^+/-^/Smad4^flox/flox^, shows reduced activation in anterior (white arrowheads) and posterior MTL (Red arrowheads) in AP-2εCre/Smad4cKO (Control n=6, cKO n=5). Statistical analysis by two tailed unpaired, t-test.

VSNs form synapses with dendrites of second order neurons within organized glomeruli of the AOB. Basal VSNs form synapses in the posterior region of the AOB (pAOB), while apical neurons project to the anterior region of the AOB (aAOB) (Brignall and Cloutier, 2015; Prince et al., 2013; Prince et al., 2009). To understand if Smad4 signaling is important for axonal organization, we performed immunostaining against OMP, Robo2, and the presynaptic marker Vesicular Glutamate Transporter 2 (VGlut2). Even though these results showed well defined glomeruli in the anterior and posterior AOB of control animals, we detected significantly fewer and larger glomeruli in the posterior portion of the AOB of AP-2εCre^+/-^/Smad4^flox/flox^ (Fig 5 C,D,F,G).

Neuronal loss in the basal VNO in adult mice and the abnormal number and size of the glomeruli suggest that Smad4 ablation in maturing VSNs compromises the long-term homeostasis of VSNs and their axonal convergence to specific glomeruli (Prince et al., 2013). To test the function of the synaptic connections between VSNs and mitral cells in the AOB, we analyzed the anterior and posterior mitral/tufted cell layer (MTL) of male animals after male whole urine exposure. By performing immunostaining against cFos, we found significant reduction in the urine exposure activated mitral cells of the posterior AOB of the AP-2εCre^+/-^/Smad4^flox/flox^, however we also detected a reduction in cFos activation in the mitral cells in the anterior AOB (Fig.5 J,K,L).

These data suggest that Smad4 ablation in basal VSNs compromises VSNs homeostasis, but does not prevent the formation of functional synapses. However, the reduction in cFos activation in the anterior AOB likely indicates a postsynaptic defect secondary to Smad4 ablation in the mitral cells of the aAOB. In fact, we previously reported that some of the mitral cells of the AOB are positive for AP-2ε lineage (Lin et al., 2018).

### Smad4 ablation in mature VSNs

We then sought to determine if Smad4 dependent BMP/TGF-β signaling is only required in a defined developmental window or if it necessary to maintain long term synaptic function and VSNs homeostasis. So, we exploited the OMPCre mouse line to conditionally ablate Smad4 in both apical and basal mature VSNs. In controls, Smad4 immunolabeling at P60 confirmed expression throughout the epithelium, while we found almost complete Smad4 ablation in both apical and basal territories after OMPCre mediated recombination (Fig 6. A,B,C). In the epithelium, the only cells positive for Smad4 are immature neurons which are strongly immunoreactive for GAP43 and negative for OMP (data not shown).

**Fig. 6.**
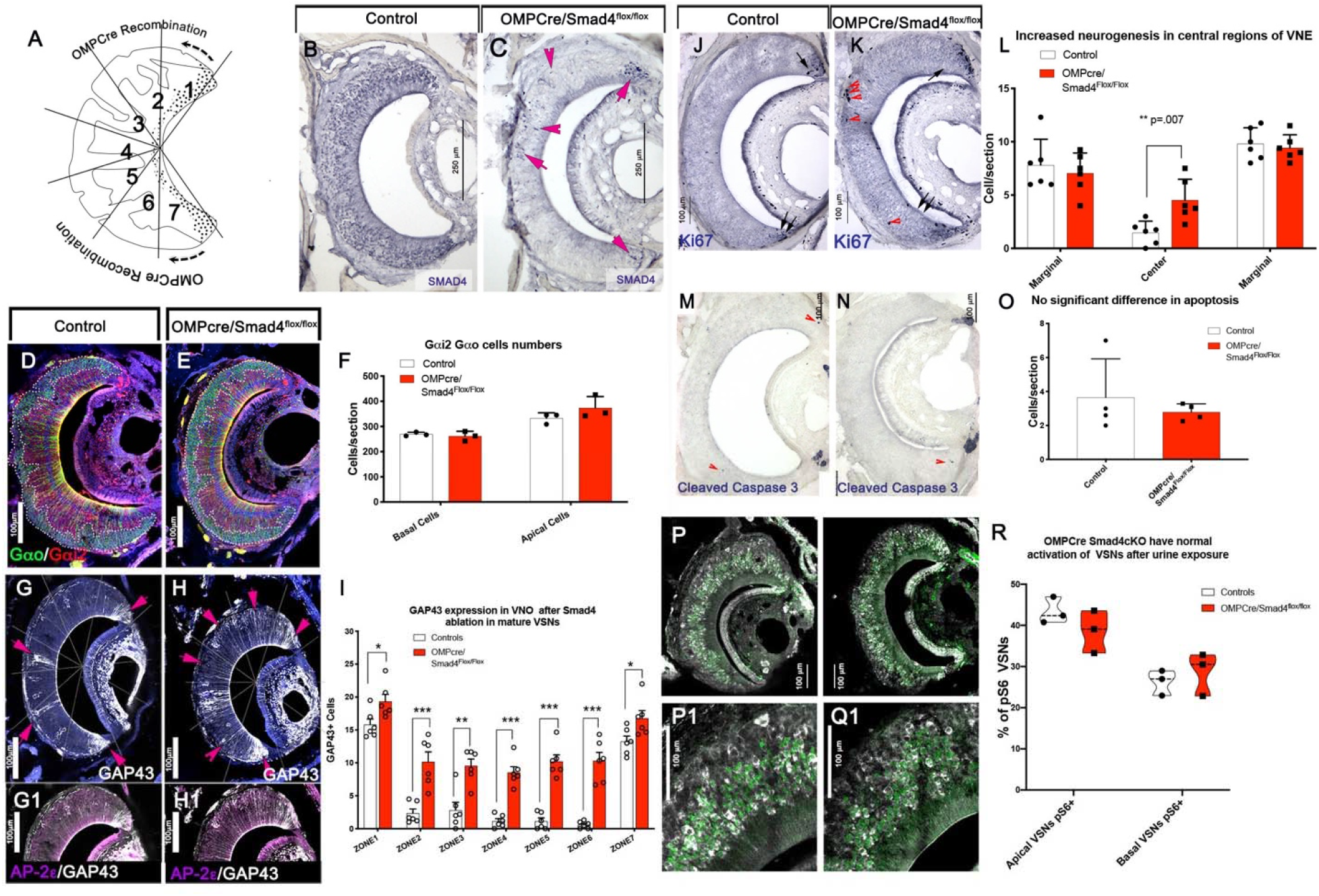
Increased GAP43 expression in VNE after Smad4 ablation in mature VSNs. A) Cartoon illustrating where OMP is expressed and therefore the Cre recombination is occurring. B,C) Smad4 ablation was confirmed by immunohistochemistry against Smad4 in control (B) and cKO (C). Few cells Smad4 positive cells remain in the cKO (arrows) as they were OMP negative. D,E,F) Immunostaining and quantification for basal marker Gαo (green) and apical marker Gαi2 (red). Quantification (F) shows no change in number of Gαo and Gαi2 VSNs in cKO as compared to control (n=3). G,G1,H,H1) Immunostaining for immature VSN marker GAP43 in control (G) and in cKO(H). Arrows indicate GAP43 immunoreactivity in VSNs from the medial zones (Zones 2-6) of the VNE. G1,H1) Immunostaining for basal specific marker AP-2ε and GAP43 indicate that GAP43 positive VSNs are predominantly AP-2ε positive. I) Quantification of GAP43 positive VSNs in control and cKO indicates significantly increased number of gap43 positive VSNs in cKO as compared to control(n=6). J,K,L) immunostaining and quantification for proliferative cell marker Ki67 in control (J) and cKO (K). Arrows indicated Ki67 positive cells in central regions of VNE. L) Quantification indicates significantly increased neurogenesis in medial zones of VNE(n=6). M,N,O) Immunostaining and quantification for cleaved caspase3 in control (M) and cKO (N) shows no change in cell death through apoptosis (n=4). P,Q) Immunostaining for pS6 and apical VSN marker Meis2 on control (P, magnification P1) and cKO (Q, magnification Q1). R) Quantification of percentage of pS6 positive apical and basal VSN(n=3). No significant changes were observed for either of the populations. Statistical analysis by two tailed unpaired, t-test.

Gross morphological observation of the VNE of OMPCre^+/-^/Smad4^flox/flox^ at P60 indicated that the localization of apical and basal VSNs within the VNO was not altered after Smad4 ablation. Further, quantification of cells positive for apical and basal markers (Gαi2 and Gαo) did not identify differences in cell composition of the VNO among genotypes (Fig. 6 D, E, F). However, we detected increased GAP43 immunoreactivity (Fig. 6 G,H,I) in the central and basal regions of the VNE of the OMPCre^+/-^/Smad4^flox/flox^ mice. This result is consistent with our results in AP-2εCre^+/-^/Smad4^flox/flox^. We also observed a significant increase in the number of Ki67+ cells in central regions of the VNE in OMPCre^+/-^/Smad4^flox/flox^ mice compared to controls (Fig. 6 J,K,L). However, we observed no changes in apoptosis among the genotypes (Fig. 6 M,N,O).

### Smad4 ablation in mature VSNs does not affect neuronal activation after odor exposure

We then examined whether Smad4 is required for neuronal activity of basal VSNs in a developmental time dependent manner. We examined VSNs neuronal activation following exposure to male urine. We used phosphorylated ribosomal subunit pS6 as a readout for VNE activation. We found no difference in the percentage of basal or apical VSNs activated between control and OMPCre^+/-^/Smad4^flox/flox^ mice (Fig. 6 P,Q,R). This result suggests that Smad4 is not required for neuronal activity after basal VSNs have reached maturity. Next, we tested whether this conditional knockout showed impaired functional connectivity of VSNs. We performed immunostaining for cFos on AOB sections of Smad4^flox/flox^ control and OMPCre^+/-^/Smad4^flox/flox^ mice exposed to male whole urine. We found no significant difference in the number of cFos positive mitral cells for either posterior or anterior AOB.

### Ablation of Smad4 in mature VSNs produces a disorganized glomerular layer in the posterior AOB but not the anterior AOB

Our analysis of VSNs’ projections to the AOB revealed that the number of glomeruli in the posterior region of AOB was significantly reduced in OMPCre^+/-^/Smad4^flox/flox^ mice compared to controls. Within the posterior AOB region, we found a significant decrease in percentage of glomeruli with area less than 200(μm)^2^ and an increase in the percentage of glomeruli with area more than 800(μm)^2^ in OMPCre^+/-^/Smad4^flox/flox^ compared to controls (Fig. 7 A-F). However, we did not observe any significant changes in average size of the glomeruli in the anterior region of AOB in OMPCre^+/-^/Smad4^flox/flox^. These data indicate that Smad4 is necessary to form the correct somatosensory map between the basal VSNs and the posterior AOB but not for apical neurons.

**Fig. 7.**
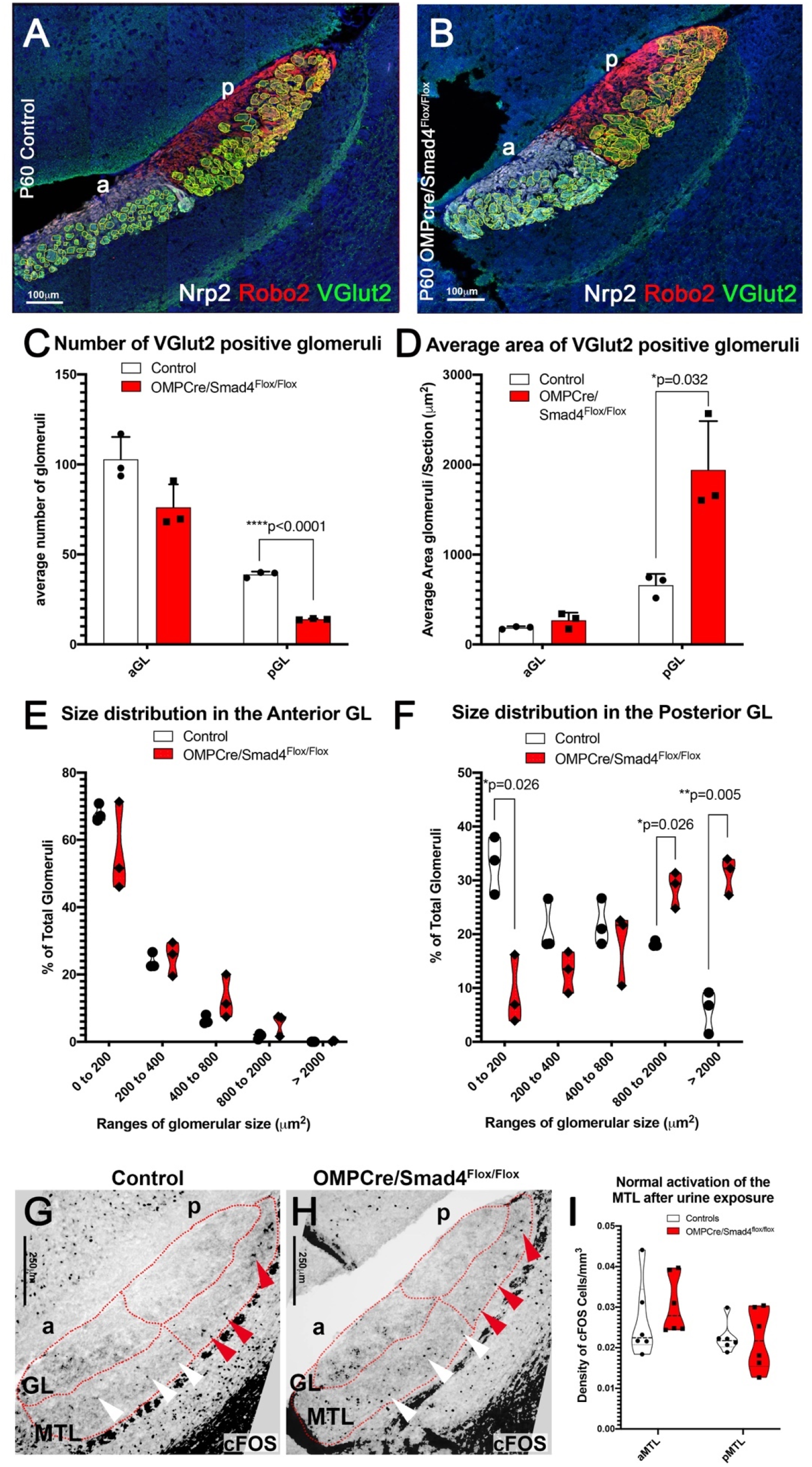
Only Basal VSNs form aberrant glomeruli at the pAOB in OMPCre/smad4cKO. A, B) Immunofluorescent staining against Nrp2,Robo2 and VGlut2 highlighting fibers projecting to the glomeruli in the aAOB and pAOB of controls and conditional mutants. C) Number of VGlut2 positive glomeruli were quantified in posterior and anterior glomerular region. Graph shows significant decrease in number of glomeruli only in posterior AOB(n=3). (D) The area of VGlut2 positive glomeruli were quantified and the graph shows a significant increase in the size of glomeruli only in the posterior AOB(n=3). (E,F) Percentage of number of glomeruli in area ranges was quantified between genotypes for anterior and posterior AOB. Graphs indicate no change in aAOB (E) and significant reduction in number of smaller glomeruli however significant increase in larger glomeruli in pAOB. G,H) Immunostaining and quantification (I) for cFOS on P60 control (Smad4^flox/flox^) and P60 OMPCre^+/-^/Smad4^flox/flox^, shows normal activation in MTL in OMPCre/Smad4cKO (n=4). Statistical analysis by two tailed unpaired, t-test.

### Loss of Smad4 alters the expression of glycoprotein and presynaptic components

We then sought to elucidate the underlying molecular mechanisms that induce aberrant glomeruli formation in Smad4cKOs in both the models. So, we performed mRNA-seq transcriptome analysis of VNO from AP-2εCre^+/-^/Smad4^flox/flox^, OMPCre+/-/Smad4^flox/flox^, relative Cre heterozygous and Smad4^flox/flox^ controls (GSE134492) mice. Our data indicated that a lack of functional Smad4 in immature (AP-2εCre^+/-^/Smad4^flox/flox)^ basal VSNs induced a significant increase in the expression of genes involved in synapse formation and glutamate release, such as Unc13-c, Synatoptagnmin-XIII, Dlg2, Adcy3, Plcb2, and Nrxn1. We confirmed the changes in Unc13-c and Nrxn1 expression by qRT-PCR (Fig. 8 E,F). While Nrxn1 levels were significantly higher in AP-2εCre^+/-^ /Smad4^flox/flox^ with respect to Smad4^flox/flox^ controls, the increase was not significant compared to AP-2εCre^+/-^ controls. However, AP-2εCre null mice (Lin et al., 2018) did not show significant changes in Nrxn1 expression (GSE110083).

**Fig. 8.**
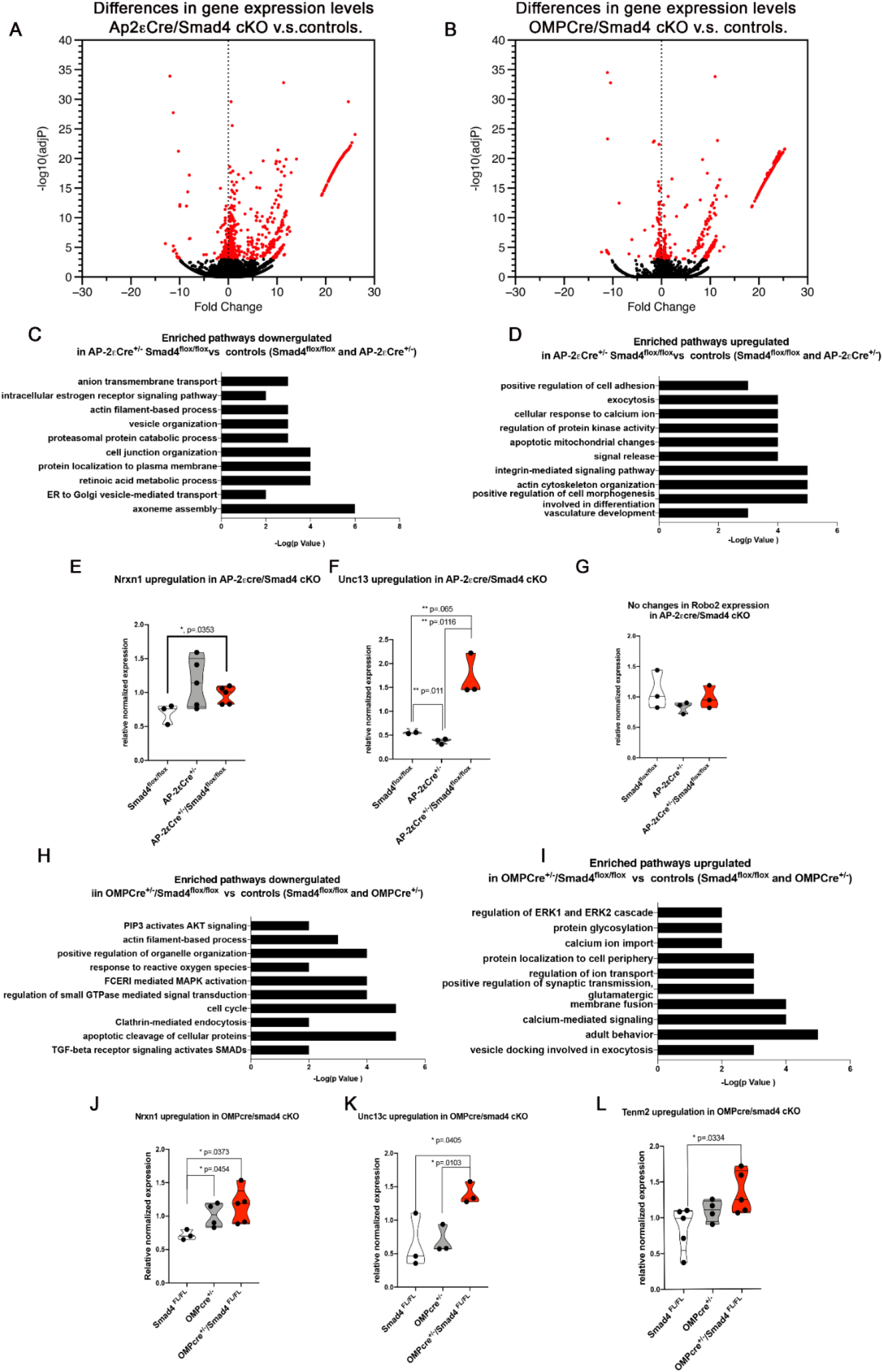
Transcriptome analysis of conditional Smad4 cKO. **A,B)** Volcano plot highlighting the changes in gene expression in the two cKO with respective controls. C,D) Enriched pathways up and downregulated in AP-2eSmad4 cKO. E,F,G) qPCR validation indicated upregulation of Nrxn1,Unc13c and no changes in Robo2. H,I) Enriched pathway up and downregulated in OMPCre/Smad4cKO. J,K,L) qPCR validation confirmed the upregulation of Nrxn1, Unc13c and Tenm2.

Transcriptome data from OMPCre^+/-^/Smad4^flox/flox^ compared to relative controls (GSE134492) also showed significant upregulation of presynaptic molecules, such as synaptic signaling (Unc13-c) and synaptic assembly (Nrxn1) (Fig.8 J,K,l). These mutants also showed upregulated tenurin-m2 (Tenm2), which participates in defining neuronal diversity and glomeruli formation (Berns et al., 2018; Mosca et al., 2012). OMP levels can also alter normal olfactory glomerular map (Albeanu et al., 2018). Our qRT-PCR analysis revealed similar changes in Nrxn1 levels between OMPCre^+/-^ and OMPCre^+/-^/Smad4^flox/flox^. These data suggest that at least part of the aberrant Nrxn1 expression observed in the OMPCre conditional KOs may partially stem from the decreased expression of OMP alone (Albeanu et al., 2018). We also observed that OMPCre^+/-^ mice had intermediate, but not significantly different, Tenm2 expression level between WT and OMPCre^+/-^/Smad4^flox/flox^ suggesting that OMP may act as a modifier modulating Tenm2 levels. However, there was a significant increase in Tenm2 expression level in OMPCre^+/-^/Smad4^flox/flox^ as compared to Smad4^flox/flox^ controls. Glomeruli of Smad4 null mutants resembles that described after loss of Kirrel family molecules (Prince et al., 2009), although our data did not indicate significant changes in Kirrel family molecules (data not shown).

## Discussion

In this study, we conditionally ablated Smad4 at two developmental time points of the VSNs in order to determine the role of Smad4 mediated TGF-β/BMP signaling in the vomeronasal organ of mice. We found that Smad4 loss-of-function in maturing basal neurons alters the homeostasis of vomeronasal epithelium leading to progressive loss of the basal population. It also reduces the activation of the surviving basal VSNs in response to male urine and leads to aberrant glomeruli formation in the pAOB. However, interference with Smad4-mediated signaling in mature VSNs does not affect neuronal activity nor survival, but causes aberrant glomeruli formation in the pAOB, but not in the aAOB. Our data suggests that Smad4-mediated signaling may play diverse role depending on the developmental maturity of the basal VSNs. We also demonstrate that TGF-β/BMP signaling may play a more critical role in homeostasis, neuronal activation and axonal convergence of basal but not apical VSNs.

Using transcriptome data, we show that the VNO expresses multiple members of the TGF-β /BMP family. We confirmed the expression of BMP4 and 6 by VSNs via in situ hybridization. Bone morphogenic proteins control proliferation, differentiation patterning, and survival of multiple tissues during embryonic development (Beites et al., 2009; Kuschel et al., 2003; Le Dreau et al., 2012; Liem et al., 2000; Liem et al., 1995; Lim et al., 2000; Schmidt et al., 1995; Tadokoro et al., 2016; Timmer et al., 2002). Even though the roles of TGF-β/BMP inductive signals during embryonic development and their role in neuronal differentiation are well-studied, their role in homeostasis and connectivity during postnatal life remains largely unexplored. Mammalian olfactory epithelia have a large neuronal diversity and undergo continuous neurogenesis (Kondo et al., 2010). These features make the olfactory system an interesting model to study the role of morphogenic signals after birth.

BMPs bind with various degrees of affinity to several ECM components, including laminins and collagens that sustain BMP signaling by binding and activating bone morphogenetic proteins. Sources of collagen IV can define a BMP signaling gradient by restricting the signaling range by sequestrating BMPs (Bunt et al., 2010; Wang et al., 2008) and facilitating its interaction with its receptors. We identified pronounced basal to apical BMP signaling gradients in the vomeronasal epithelium that were independent of gradients in BMP expression.

By generating AP-2εCre/Smad4 and OMPCre/Smad4 conditional mouse mutants, we experimentally tested the role BMP and TGF-β Smad4 dependent intracellular signaling in maturing basal VSNs (AP-2εCre/Smad4) and in mature apical and basal VSNs (OMPCre/Smad4). Together results from both models indicate that Smad4 intracellular signaling is necessary for basal VSNs survival and function in a certain developmental period, but our data suggests that it is dispensable for apical VSNs. Ablating Smad4 signaling in different maturation stages of VSNs exerts different outcomes. By ablating Smad4 in maturing basal VSNs, we found that Smad4 loss-of-function is compatible with the formation of the vomeronasal epithelium, as we could not distinguish the VNO of mutant animals from controls at two weeks after birth. However, we found compromised functionality of basal VSNs, aberrant connectivity to the accessory olfactory and reduced basal VSNs’ neuronal survival in the vomeronasal system in adult P60 animals.

By analyzing cFos activation in the AOBs of AP-2εCre/Smad4 mutants after male urine exposure we detected reduced activation of the mitral cells in both anterior and posterior aAOB. While the reduction of cFOS in the posterior AOB likely reflects presynaptic defects secondary to Smad4 ablation in the basal VSNs, the reduction in cFOS activation in the aAOB likely reflect a post-synaptic defect. In fact, we previously reported that the mitral cells of the aAOB but not the ones of the pAOB are positive for AP2e lineage. Reduced cFOS activation in the aAOB of AP-2εCre/Smad4 mutants suggests a role for Smad4 in controlling maturation or function of the anterior mitral cells.

Conversely, Smad4 ablation in mature (OMP positive) VSNs did not affect the function or maintenance of basal VSNs, but affected normal formation of glomeruli in the posterior accessory olfactory bulb as observed in AP-2εCreSmad4cko. In both mutants, we found less and abnormally larger glomeruli only in the posterior AOB.

Retrograde BMP signaling in neurons can control synaptic growth at neuromuscular junctions and define the neuronal identity and somatosensory map formation of the trigeminal nerve in rodents (Ball et al., 2010; Banerjee and Riordan, 2018; Berke et al., 2013; Fuentes-Medel and Budnik, 2010; Hegarty et al., 2013; Hodge et al., 2007; Liao et al., 2018; Piccioli and Littleton, 2014). Publicly available gene expression atlases indicate that the olfactory bulbs express various BMPs, including |BMP 1,15, 6 and 8B (Allen brain atlas-ISH data). Our experimental approach does not permit dissociating aberrant local vomeronasal BMPs signaling and putative retrograde BMP signaling from the AOBs. We acknowledge our observations may reflect compromised local and retrograde morphogenic signaling. However, we feel this is unlikely considering we saw similar pattern of p-Smad 1/5/8 in VNE of Arx-1 Null mice, which lack olfactory bulbs, as the controls (Data not shown).

Using OMPCre mediated Smad4 ablations to alter both apical and basal VSNs, we observed significant changes in basal VSNs projections only. We found that the basal VSNs actively transduce higher levels of BMP signaling. We speculate that the basal to apical gradients we observed in the vomeronasal epithelium reflect local morphogenic signaling.

Selective connectivity of VSNs to the mitral cells and the formation of glomeruli in the accessory olfactory bulbs is defined by expression of specific vomeronasal receptors, guidance cues like Slit and Semaphorins, and adhesion molecules, such as the Kirrel family (Belluscio et al., 1999; Brignall and Cloutier, 2015; Cloutier et al., 2002; Del Punta et al., 2000; Ishii and Mombaerts, 2011; Prince et al., 2013; Prince et al., 2009). After Smad4 ablation in maturing and in mature VSNs, we observed an aberrant number and size of glomeruli in the posterior AOB. In AP-2εCre Smad 4 cKO, we found reduced neuronal activity in the vomeronasal epithelium, but not in the OMPCreSmad4cKO mutants. The glomerular morphology in AP-2eCre Smad4cKOs resembles that described in Kirrel 2,3 knockout mice (Prince et al., 2013, Brignall et al 2018). Activity dependent levels of cAMP (Prince et al., 2013, Albeanu et al., 2018) can modulate expression of Kirrel 2,3 and refine glomeruli formation. We found similar aberrant glomeruli in both Smad4cKO models, even though neuronal activity was hampered only in AP-2εSmad4cKO mutants. Based on these data, we speculate that the observed phenotype may be independent from the loss of neuronal activity.

Analyzing transcriptome data of both mutants with Smad4 ^flox/flox^ and relative Cre controls, we did not find any indication of statically significant changes in expression levels of Kirrel molecules. Rather, we found upregulation of several presynaptic genes, including Nrxn1 and Unc13c (See supplementary data). One upregulated signaling protein after loss of Smad4, Nrxn1, is a transmembrane synaptic adhesive molecule that regulates the synaptic architecture and function in the brain. The interaction between trans-synaptic molecules, such as Neurexin1 and NCAM1, is crucial for defining pre and post synaptic target recognition alignment during synaptogenesis and synaptic transmission (Gerrow and El-Husseini, 2006, Scheiffele, 2003, Yamagata et al., 2003). If changes in Nrxn1 expression level (Ko et al., 2009; Sudhof, 2017) define synaptic identity, we suggest that compatibility and selective connectivity warrants future investigation.

In OMPCre^+/-^/Smad4^flox/flox^ mutants, we also found significant upregulation of the glycoprotein Tenm2, a molecule crucial for defining synaptic matching and glomeruli formation of olfactory neurons in drosophila (Hong et al., 2012). In both Smad4 conditional models, we observed increased immunoreactivity for GAP43, which suggests compromised/incomplete neuronal maturation. We propose that Smad4 mediated signaling may modulate the required gene expression and chromatin rearrangements to complete VSNs maturation. Proteins of the Nrxn1 family undergo dynamic expression as neurons mature during development (Harkin et al., 2017). Based on our findings in OMPCre^+/-^/Smad4^flox/flox^ (cKO) mutants, we speculate that changes in Nrx1 expression arise from an overall compromised maturation rather than a direct Smad4 dependent regulation. Though both models showed changes in the expression of multiple genes that contribute to synaptic formation and maturation, we did not detect obvious loss-of-function for neuronal connections between VSNs and the brain. In fact, both AP-2εCre and OMPCre Smad4cKOs mutants could activate mitral cells in the brain after urine exposure.

Overall, we found that Smad4 signaling is necessary for connectivity and survival in maturing VSNs. While Smad4 can lead to formation of abnormal glomeruli, it does not affect survival of mature VSNs. By ablating Smad4 in maturing or in mature VSNs, we revealed dynamic requirements for Smad4 signaling. Based on the phenotypes we observed, we identified a bias in the requirement of Smad4 mediated transcription for basal compared to apical VSNs. We speculate that the relative position of VSNs with respect to BMP signaling in the epithelium may define the expression level of genes for glycoproteins, adhesion molecules and presynaptic components that then modulate the specificity and accuracy of synapse formation in the AOB.

## Material and Methods

### Animals

AP-2εCre line (TfAP-2e^tm1(cre)Will^) was donated by Dr. Trevor Williams (Department of Craniofacial Biology, University of Colorado). OMPCre line was donated by Dr Paul Feinstein (Hunter college, City University of New York). Smad4 ^flox/flox^ (*Smad4^tm2.1Cxd^*/J, Stock # 017462) and R26RYFP (B6.129X1-*Gt(ROSA)26Sor^tm1(EYFP)Cos^*/J, Stock #00614) were purchased from Jackson Laboratories. Experimental analyses were carried out in offspring both homozygous for floxed Smad4 and AP-2εCre^+/-^ (AP-2εCre^+/-^ /Smad4^flox/flox^ or AP-2εCre^+/-^/Smad4cKO) or Smad4^flox/flox^ and OMPCre^+/-^(OMPCre^+/-^/ Smad4^flox/flox^ or OMPCre^+/-^/ Smad4cKO), hence-forward referred to as conditional Smad4 mutants. The controls used were Smad4^flox/flox^ unless otherwise indicated. These mice appeared healthy and survived to adulthood. Mice of either sex were used for ISH and IHC experiments. All experiments involving mice were approved by the University at Albany Institutional Animal Care and Use Committee (IACUC).

### Tissue Preparation

Tissue was collected after transcardial perfusion with first PBS and then 3.7% formaldehyde in PBS. Mouse brains were additionally immersion-fixed in 1% formaldehyde at 4°C overnight. Noses were immersion fixed in 3.7% formaldehyde in PBS at 4°C overnight, decalcified in 500mM EDTA for 3-7 days. After fixation and decalcification samples were cryoprotected in 30% sucrose overnight at 4°C, then embedded in O.C.T. (Tissue-TeK) and stored at −80°C.

Samples were cryosectioned using CM3050S Leica cryostat at 16μm (nose) and 20 μm (brain) collected on Superfrost Plus Micro Slides (VWR). All sections were stored at −80°C until ready for staining or in situ hybridization.

### Immunohistochemistry

Primary antibodies and concentrations used in this study were, *Gt α-AP-2ε(2µg/ml,sc-131393 X, Santa Cruz) *Rb α-Cleaved Caspase-3 (1:1000,AB3623,Millipore), *Ms α-Gαi2 (1:200, 05-1403, Millipore), Rb α-Gαo(1:1000, 551, Millipore), Rb α-GAP43 (1:500, 16053, abcam), *Rb α-Ki67 (1:1000, AB9260,Millipore), Gt α-NPN2 (1:4000, AF 567,R&D Systems), Gt α-OMP (1:4000, 5441001, WAKO), *Ms α-Robo2 (1:100, 376177, Santa Cruz), Rb α-V2R2 (1:4000, Gift from Dr. Roberto Tirindelli, Univ. di Parma, Italy), *Rb α-pSmad 2 (1:800, AB 3849, Millipore), *Rb α-pSmad1,5,8 (1:50, AB3848-I, Millipore), Rb α-Smad4 (1:200, AB40759, Abcam), Chk α-VGlut2(1:500, 135416, Synaptic Systems), *Rbα-pS6(ser 240/244) (1:500, D68F8,cell signaling tech), *Rbα-cFOS (1:250, cell signaling tech). Microwave antigen retrieval in citrate buffer pH 6 was performed for the indicated antibodies (*)(Forni et al., 2006). All primary antibodies were incubated at 4°C over-night.

For immunoperoxidase staining procedures, slides were processed using standard protocols (Forni et al., 2013) and staining was visualized (Vectastain ABC Kit, Vector, Burlingame, CA) using diaminobenzidine (DAB) in a glucose solution containing glucose oxidase to generate hydrogen peroxide. After immunostaining sections were counterstained with methyl green.

For immunofluorescence, species-appropriate secondary antibodies were conjugated with Alexa-488, Alexa-594, or Alexa-568 (Molecular Probes). Immunofluorescent sections were counterstained with 4′,6′-diamidino-2-phenylindole (DAPI, 1:3000; Sigma-Aldrich) and mounted with FluoroGel (Electron Microscopy Services). Confocal microscopy pictures were taken on a Zeiss LSM 710 microscope. Bright field and epifluorescence pictures were taken on a Leica DM4000 B LED fluorescence microscope attached to Leica DFC310 FX camera. Images were analyzed and quantified using FIJI/ImageJ software.

### In Situ Hybridization

Digoxigenin-labeled cRNA probes against Smad4 Exon 8, BMP4 and 6 were prepared by *in vitro* transcription (DIG RNA labeling kit; Roche Diagnostics). Plasmids to generate cRNA probe for Smad4 was gifted by Prof. Dr. Rolf Zeller, University of Basel, Switzerland and for BMP4 and 6 was provided by Dr. Kapil Bharti, Ocular and Stem Cell Translational Research Unit, NEI. In situ hybridization was performed as described (Lin etal., 2018) and visualized by immunostaining with an alkaline phosphatase conjugated anti-DIG (1:1000), and NBT/BCIP developer solution (Roche Diagnostics).

### Male whole urine exposure

2-month-old male Smad4^flox/flox^ control and conditional KO animals were single caged 2-3 days before the experiment. Male urine was collected and pooled from mice of mixed background. Male control and experimental mice were exposed to male urine and then perfused and collected 90 mins after exposure. The brains were sectioned parasagittal and the noses were sectioned coronal and immunostained for cFos and pS6 respectively. BS-Lectin I conjugated to Rhodamine (Vector) was used in order to distinguish between anterior and posterior AOB (Prince et al., 2013).

### Quantification and statistical analyses of microscopy data

Data was collected from mice kept in same housing conditions. Measurements of areas, thickness and cell counts were performed on digital pictures using FIJI/ImageJ. For some of the antigens that have different expression from the tips of the VNO to the center of the epithelium (where cells are mostly mature neurons) cell counts were done overlapping a mask to equally divide the VNO in seven 36-degree angled zones (Rosa-Prieto de la et al., 2010).

For all AOB quantifications most medial 3 parasagittal sections per AOB were used. We used confocal images of triple staining against VGlut2, Npn2, and Robo2 to trace the VGlut2 positive glomeruli in both Npn2 positive anterior and Robo2 positive posterior AOB. Using Fiji imaging tool, we generated maximum intensity projection image from the 3D stack to increase signal to noise ratio and manually traced each glomerulus. We analyzed the average number and glomeruli area in anterior and posterior AOB for each section. This was then used to determine the overall average of all technical replicates from each biological replicate. Number of glomeruli within area ranges was calculated and then compared between genotypes.

In animals ≥P15, the most medial 6-8 VNE sections were quantified for each series and averaged. Data form each genotype was grouped and used for statistical analysis. Statistical analysis was performed using GraphPad, Prism 7.0b. Two tailed unpaired t-test were used for statistical analyses and p-value less than 0.05 was considered statistically significant. Sample sizes and p-Values are indicated in figure legends or in each graph.

### RT-PCR

cDNA was synthesized from 1000 ng of RNA extracted from whole VNO of P21 Smad4^flox/flox^ animals (n=3) by using the kit by Invitrogen Super ScriptIII First Strand (Cat# 18080400). Semi-quantitative RT-PCR for BMP2,3,4,6,7, TGF-β1, TGF-β2 and GDF10 was done using cDNA generated from RNA isolated from dissected vomeronasal organ of Smad4^flox/flox^ mice using the primers indicated in Supplementary Table 1.

### qPCR

cDNA was synthesized from 500 ng of RNA extracted from whole VNO of control animals (P21) (Smad4^flox/flox^ and AP-2εCre^+/-^ or Smad4^flox/flox^ and OMPCre^+/-^) and (P21) cKOs (AP-2εCre^+/-^;Smad4^flox/flox^ or OMPCre^+/-^;Smad4^flox/flox^) by using the kit by Invitrogen Super Script III first strand catalogue # 18080400. The expression of Nrxn1, Unc13c, Tenm2 and Robo2 was validated using qRT PCR using primers listed in Supplementary Table 2. Three technical replicates of each sample were run and a standard curve was used to get relative quantities, which were then normalized to ubiquitin C.

### Transcriptome analysis

Total RNA was isolated using PureLink RNA mini kit Cat # 12183018A from dissected vomeronasal organ of control (Smad4^flox/flox^, AP-2εCre^+/-^ and OMPCre^+/-^) and Smad4cKOs (AP-2εCre^+/-^/Smad4^flox/flox^ and OMPCre^+/-^/Smad4^flox/Flox^). RNA with RIN number higher than 8 on Agilent Bioanalyzer was used for further experiments. Libraries for mRNA seq were prepared using NEXTflex Rapid directional mRNA-seq Bundle, cat# 5138-10). To prepare libraries, PolyA+ RNA was isolated from 1 microgram of total RNA using 2 rounds of selection with poly dT magnetic beads (BioO Scientific) and used as the template for 1^st^ strand synthesis using random hexamers. Second strand synthesis was performed using the standard RNAseH-mediated nicking with dUTP for preserving strandedness. Double stranded cDNA was then used as the template for library construction following the manufacturer’s recommendations for 1 microgram of starting total RNA (BioO Scientific NextFlex). Resulting libraries were quantified using the NEBNext Library Quant Kit for Illumina (New England Biolabs) and sizes were confirmed using an Agilent Bioanalyzer. Libraries were sequenced on an Illumina NextSeq 500 instrument for 75 cycles in single-read orientation. The resulting raw sequencing files were quantified using salmon (v 0.14.0) in quant mode and the – validateMappings flag using the ENSMBL mm10 GRC38 mouse CDS assembly. Raw read counts per gene were imported into R and differential gene expression was determined using DESeq2 (v 1.22.2). Gene Ontology was performed using the MetaScape package.

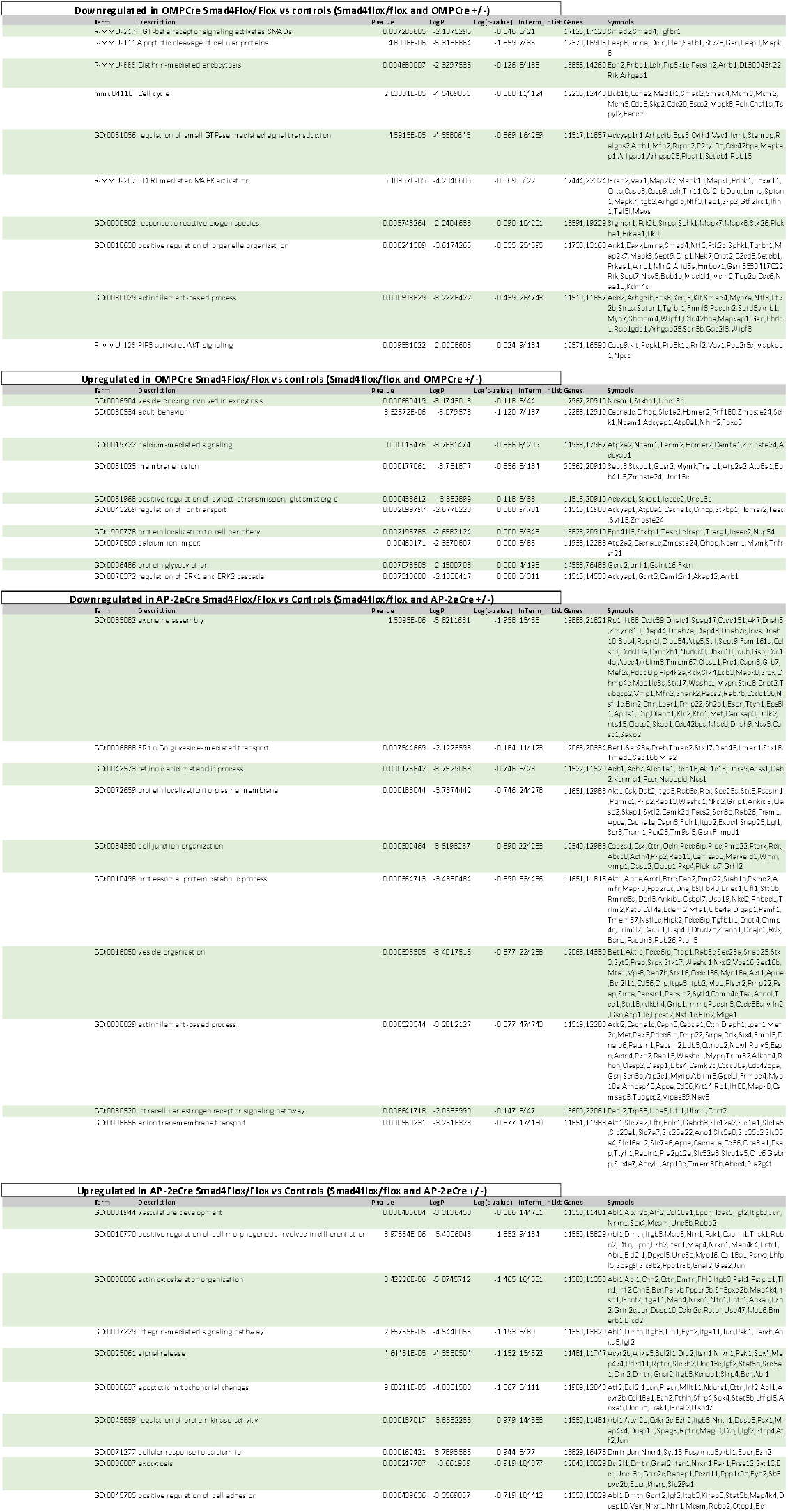

**Table 1.**
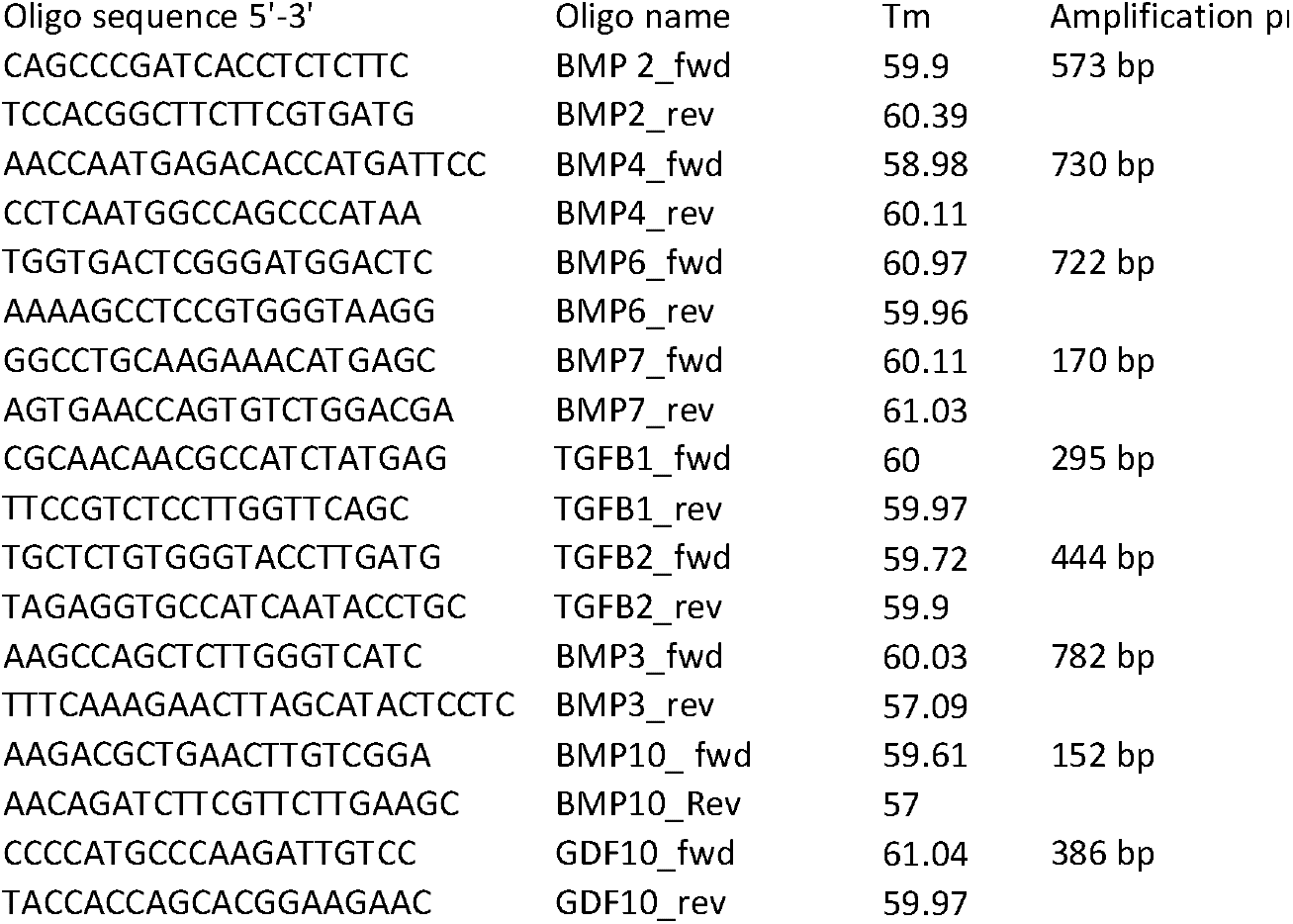
Primers used for semi quantitative RT-PCR

**Table 1.**
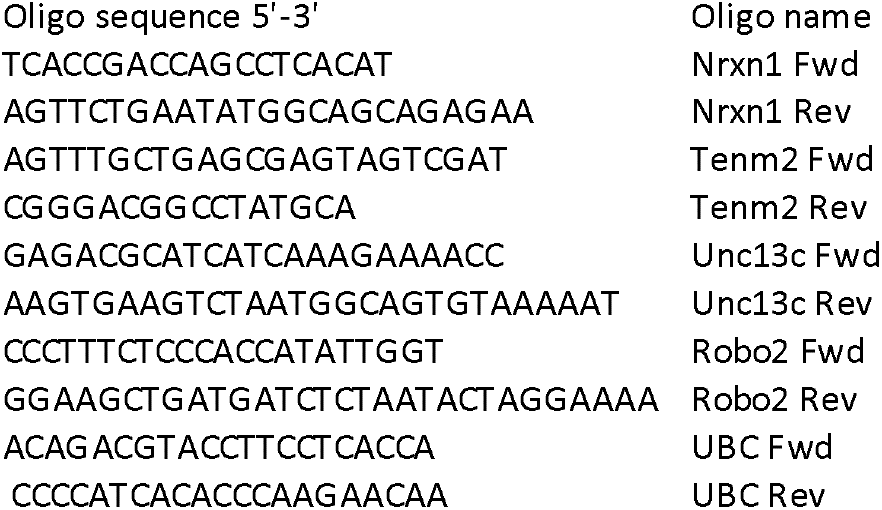
Primers used for qPCR

## Acknowledgments

Research reported in this publication was supported by the Eunice Kennedy Shriver National Institute of Child Health and Human Development of the National Institutes of Health under the Award Numbers 1R01HD097331-01 (PF) and by the National Institute of Deafness and other Communication Disorders of the National Institutes of Health under Award Number 1R01DC017149-01A1 (PF). The content of this manuscript is solely the responsibility of the authors and does not necessarily represent the official views of the National Institutes of Health.

---

**S1:**
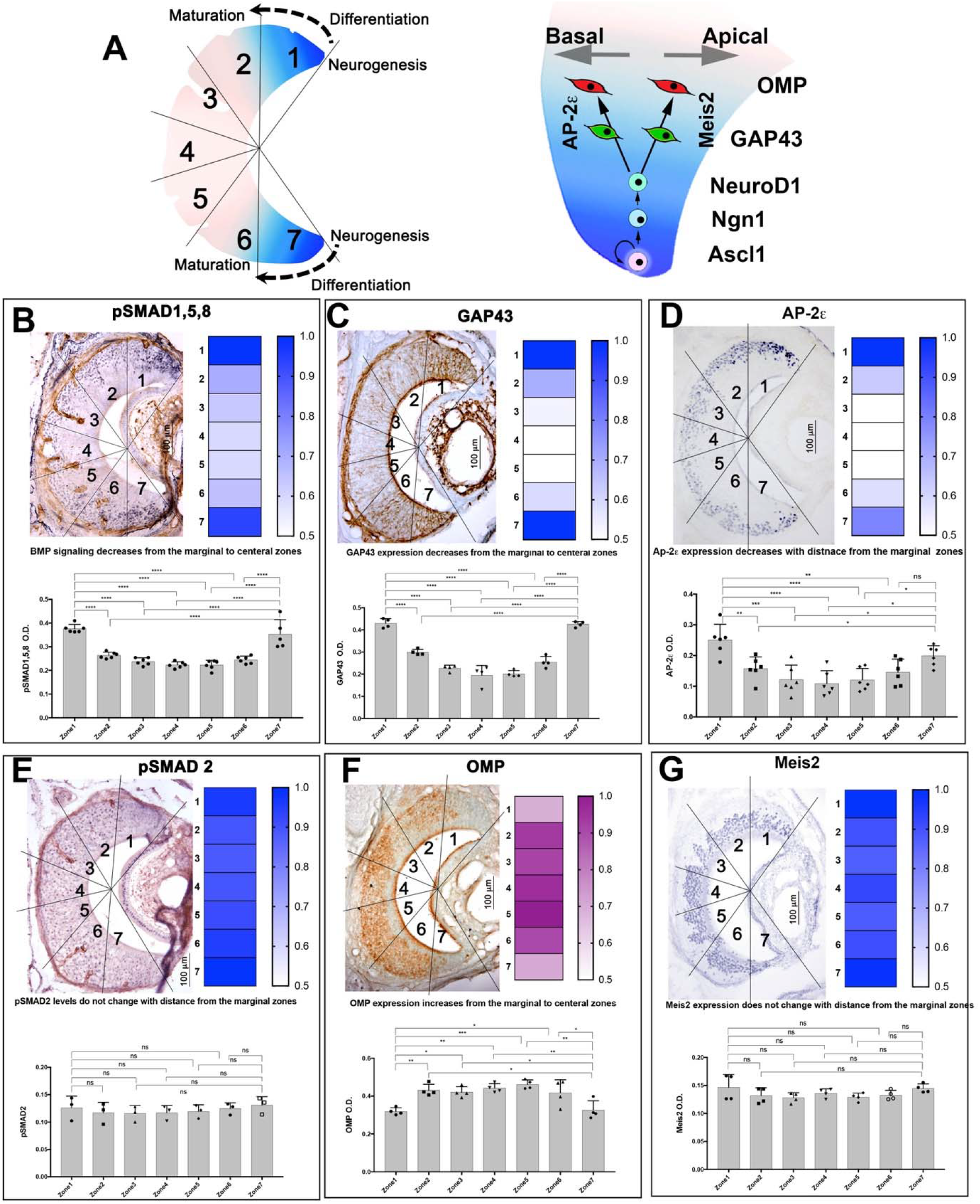
Spatial correlation between ability to transduce BMP, AP-2ε expression, and GAP43 expression. **A)** Cartoon illustrating neurogenesis, differentiation and maturation in the VNE. Most of the neurogenesis occurs at the margins followed by differentiation into apical or basal VSNs which then undergo maturation. The Ascl1 neuronal progenitor cell self-renews and divides to form first Ngn1 and then NeuroD positive precursor. These then differentiate into AP-2ε positive basal VSN or Meis2 positive apical VSN. At this point these are immature neurons and start to express GAP43. Upon reaching functional maturity they express olfactory marker protein (OMP). B) Immunohistochemistry for pSmad 1,5,8 and its densitometric analysis along the marginal, intermediate and central zones of VNE. Graph and heat map showing cells strongly positive for pSmad1,5,8 are confined to marginal zones with immunoreactivity significantly decreasing towards the central zones. C) Immunohistochemistry for immature marker GAP43 and its densitometric analysis along the VNE. Graph and heat map show immature neurons are largely present in the marginal zones of VNE. D) Immunohistochemistry for AP-2ε and its densitometric analysis. Graph and heat map show strong AP-2ε expressing cells in the marginal zones on VNE. E,F,G) Immunohistochemistry and densitometric analysis of pSmad2, OMP and Meis2. Graph and heat map show equal distribution of cells expressing these markers in all the zones of VNE. OD measured in each zone following DAB staining after dividing the VNE in 7 zones, unpaired t-Test p (*<0.05), +/-SEM, (N=3 animals;3-4 sections per animal).

**S2:**
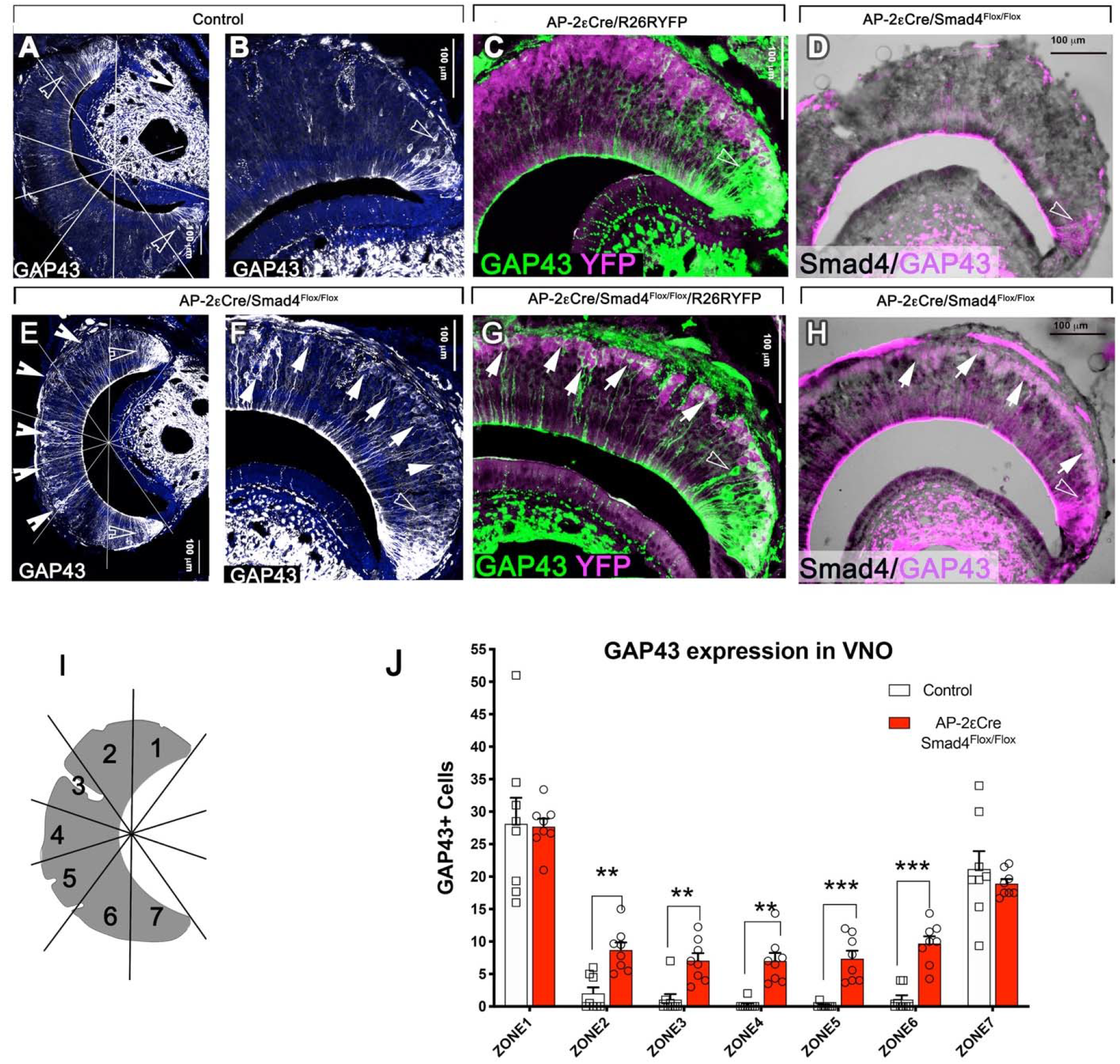
Increased GAP43 expression with early Smad4 conditional ablation. A,E) GAP43 immunostaining in control (A) and cKO (E). White notched arrows point at GAP43 positive cells in the medial regions of VNE in the cKO (E), while in control GAP43 positive cells are mostly restricted to the marginal regions, empty notched arrows(A). B,F) magnification of A and E respectively. C,G) Immunostaining for AP-2ε driven recombination (YFP, magenta) and GAP43 (green) in control (C) and cKO (G). White arrows point at tracing positive GAP43 positive cells in the medial regions of VNE in cKO, while empty arrows point at non-traced GAP43 positive cells which are restricted to marginal zones(G). D,H) Immunostaining for GAP43 (magenta) and Smad4 (black) in control (D) and cKO (H). I) Cartoon illustrating how the VNE was divided in 7 zones for quantification purposes. J) Graph representing significant increase in GAP43 positive cells in the medial regions of VNE in CKO, while marginal regions had comparable number of GAP43 positive cells in control and cKO (N=6).

## REFERENCES

Albeanu, D.F., Provost, A.C., Agarwal, P., Soucy, E.R., Zak, J.D., and Murthy, V.N. (2018). Olfactory marker protein (OMP) regulates formation and refinement of the olfactory glomerular map. Nat Commun 9, 5073.

Ball, R.W., Warren-Paquin, M., Tsurudome, K., Liao, E.H., Elazzouzi, F., Cavanagh, C., An, B.S., Wang, T.T., White, J.H., and Haghighi, A.P. (2010). Retrograde BMP signaling controls synaptic growth at the NMJ by regulating trio expression in motor neurons. Neuron 66, 536–549.

Banerjee, S., and Riordan, M. (2018). Coordinated Regulation of Axonal Microtubule Organization and Transport by Drosophila Neurexin and BMP Pathway. Sci Rep 8, 17337.

Beites, C.L., Kawauchi, S., and Calof, A.L. (2009). Olfactory Neuron Patterning and Specification. Dev Neurobiol 7, 145–156.

Belluscio, L., Koentges, G., Axel, R., and Dulac, C. (1999). A map of pheromone receptor activation in the mammalian brain. Cell 97, 209–220.

Benazet, J.D., Pignatti, E., Nugent, A., Unal, E., Laurent, F., and Zeller, R. (2012). Smad4 is required to induce digit ray primordia and to initiate the aggregation and differentiation of chondrogenic progenitors in mouse limb buds. Development 139, 4250–4260.

Berke, B., Wittnam, J., McNeill, E., Van Vactor, D.L., and Keshishian, H. (2013). Retrograde BMP signaling at the synapse: a permissive signal for synapse maturation and activity-dependent plasticity. J Neurosci 33, 17937–17950.

Berns, D.S., DeNardo, L.A., Pederick, D.T., and Luo, L. (2018). Teneurin-3 controls topographic circuit assembly in the hippocampus. Nature 554, 328–333.

Brann, J.H., and Firestein, S. (2010). Regeneration of new neurons is preserved in aged vomeronasal epithelia. J Neurosci 30, 15686–15694.

Brignall, A.C., and Cloutier, J.F. (2015). Neural map formation and sensory coding in the vomeronasal system. Cell Mol Life Sci 72, 4697–4709.

Bunt, S., Hooley, C., Hu, N., Scahill, C., Weavers, H., and Skaer, H. (2010). Hemocyte-secreted type IV collagen enhances BMP signaling to guide renal tubule morphogenesis in Drosophila. Dev Cell 19, 296–306.

Chamero, P., Leinders-Zufall, T., and Zufall, F. (2012). From genes to social communication: molecular sensing by the vomeronasal organ. Trends Neurosci 35, 597–606.

Cloutier, J.F., Giger, R.J., Koentges, G., Dulac, C., Kolodkin, A.L., and Ginty, D.D. (2002). Neuropilin-2 mediates axonal fasciculation, zonal segregation, but not axonal convergence, of primary accessory olfactory neurons. Neuron 33, 877–892.

Coleman, J.H., Lin, B., Louie, J.D., Peterson, J., Lane, R.P., and Schwob, J.E. (2019). Spatial Determination of Neuronal Diversification in the Olfactory Epithelium. J Neurosci 39, 814–832.

de la Rosa-Prieto, C., Saiz-Sanchez, D., Ubeda-Banon, I., Argandona-Palacios, L., Garcia-Munozguren, S., and Martinez-Marcos, A. (2010). Neurogenesis in subclasses of vomeronasal sensory neurons in adult mice. Dev Neurobiol 70, 961–970.

Del Punta, K., Puche, A., Adams, N.C., Rodriguez, I., and Mombaerts, P. (2002). A divergent pattern of sensory axonal projections is rendered convergent by second-order neurons in the accessory olfactory bulb. Neuron 35, 1057–1066.

Del Punta, K., Rothman, A., Rodriguez, I., and Mombaerts, P. (2000). Sequence diversity and genomic organization of vomeronasal receptor genes in the mouse. Genome Res 10, 1958–1967.

Dulac, C. (2000). Sensory coding of pheromone signals in mammals. Curr Opin Neurobiol 10, 511–518.

Enomoto, T., Ohmoto, M., Iwata, T., Uno, A., Saitou, M., Yamaguchi, T., Kominami, R., Matsumoto, I., and Hirota, J. (2011). Bcl11b/Ctip2 controls the differentiation of vomeronasal sensory neurons in mice. J Neurosci 31, 10159–10173.

Forni, P.E., Bharti, K., Flannery, E.M., Shimogori, T., and Wray, S. (2013). The indirect role of fibroblast growth factor-8 in defining neurogenic niches of the olfactory/GnRH systems. J Neurosci 33, 19620–19634.

Forni, P.E., Scuoppo, C., Imayoshi, I., Taulli, R., Dastru, W., Sala, V., Betz, U.A., Muzzi, P., Martinuzzi, D., Vercelli, A.E., et al. (2006). High levels of Cre expression in neuronal progenitors cause defects in brain development leading to microencephaly and hydrocephaly. J Neurosci 26, 9593–9602.

Fuentes-Medel, Y., and Budnik, V. (2010). Menage a Trio during BMP-Mediated Retrograde Signaling at the NMJ. Neuron 66, 473–475.

Garamszegi, N., Garamszegi, S.P., Samavarchi-Tehrani, P., Walford, E., Schneiderbauer, M.M., Wrana, J.L., and Scully, S.P. (2010). Extracellular matrix-induced transforming growth factor-beta receptor signaling dynamics. Oncogene 29, 2368–2380.

Giacobini, P., Benedetto, A., Tirindelli, R., and Fasolo, A. (2000). Proliferation and migration of receptor neurons in the vomeronasal organ of the adult mouse. Brain Res Dev Brain Res 123, 33–40.

Harkin, L.F., Lindsay, S.J., Xu, Y., Alzu’bi, A., Ferrara, A., Gullon, E.A., James, O.G., and Clowry, G.J. (2017). Neurexins 1-3 Each Have a Distinct Pattern of Expression in the Early Developing Human Cerebral Cortex. Cereb Cortex 27, 216–232.

Hegarty, S.V., O’Keeffe, G.W., and Sullivan, A.M. (2013). BMP-Smad 1/5/8 signalling in the development of the nervous system. Prog Neurobiol 109, 28–41.

Hodge, L.K., Klassen, M.P., Han, B.X., Yiu, G., Hurrell, J., Howell, A., Rousseau, G., Lemaigre, F., Tessier-Lavigne, M., and Wang, F. (2007). Retrograde BMP signaling regulates trigeminal sensory neuron identities and the formation of precise face maps. Neuron 55, 572–586.

Hong, W., Mosca, T.J., and Luo, L. (2012). Teneurins instruct synaptic partner matching in an olfactory map. Nature 484, 201–207.

Hovis, K.R., Ramnath, R., Dahlen, J.E., Romanova, A.L., LaRocca, G., Bier, M.E., and Urban, N.N. (2012). Activity regulates functional connectivity from the vomeronasal organ to the accessory olfactory bulb. J Neurosci 32, 7907–7916.

Ishii, T., and Mombaerts, P. (2011). Coordinated coexpression of two vomeronasal receptor V2R genes per neuron in the mouse. Mol Cell Neurosci 46, 397–408.

Isogai, Y., Si, S., Pont-Lezica, L., Tan, T., Kapoor, V., Murthy, V.N., and Dulac, C. (2011). Molecular organization of vomeronasal chemoreception. Nature 478, 241–245.

Ji, S.J., and Jaffrey, S.R. (2012). Intra-axonal translation of SMAD1/5/8 mediates retrograde regulation of trigeminal ganglia subtype specification. Neuron 74, 95–107.

Ko, J., Zhang, C., Arac, D., Boucard, A.A., Brunger, A.T., and Sudhof, T.C. (2009). Neuroligin-1 performs neurexin-dependent and neurexin-independent functions in synapse validation. EMBO J 28, 3244–3255.

Kondo, K., Suzukawa, K., Sakamoto, T., Watanabe, K., Kanaya, K., Ushio, M., Yamaguchi, T., Nibu, K., Kaga, K., and Yamasoba, T. (2010). Age-related changes in cell dynamics of the postnatal mouse olfactory neuroepithelium: cell proliferation, neuronal differentiation, and cell death. J Comp Neurol 518, 1962–1975.

Kuschel, S., Ruther, U., and Theil, T. (2003). A disrupted balance between Bmp/Wnt and Fgf signaling underlies the ventralization of the Gli3 mutant telencephalon. Dev Biol 260, 484–495.

Le Dreau, G., Garcia-Campmany, L., Rabadan, M.A., Ferronha, T., Tozer, S., Briscoe, J., and Marti, E. (2012). Canonical BMP7 activity is required for the generation of discrete neuronal populations in the dorsal spinal cord. Development 139, 259–268.

Lee, K.J., Dietrich, P., and Jessell, T.M. (2000). Genetic ablation reveals that the roof plate is essential for dorsal interneuron specification. Nature 403, 734–740.

Liao, E.H., Gray, L., Tsurudome, K., El-Mounzer, W., Elazzouzi, F., Baim, C., Farzin, S., Calderon, M.R., Kauwe, G., and Haghighi, A.P. (2018). Kinesin Khc-73/KIF13B modulates retrograde BMP signaling by influencing endosomal dynamics at the Drosophila neuromuscular junction. PLoS Genet 14, e1007184.

Liem, K.F., Jr., Jessell, T.M., and Briscoe, J. (2000). Regulation of the neural patterning activity of sonic hedgehog by secreted BMP inhibitors expressed by notochord and somites. Development 127, 4855–4866.

Liem, K.F., Jr., Tremml, G., Roelink, H., and Jessell, T.M. (1995). Dorsal differentiation of neural plate cells induced by BMP-mediated signals from epidermal ectoderm. Cell 82, 969–979.

Lim, D.A., Tramontin, A.D., Trevejo, J.M., Herrera, D.G., Garcia-Verdugo, J.M., and Alvarez-Buylla, A. (2000). Noggin antagonizes BMP signaling to create a niche for adult neurogenesis. Neuron 28, 713–726.

Lin, J.M., Taroc, E.Z.M., Frias, J.A., Prasad, A., Catizone, A.N., Sammons, M.A., and Forni, P.E. (2018). The transcription factor Tfap2e/AP-2epsilon plays a pivotal role in maintaining the identity of basal vomeronasal sensory neurons. Dev Biol.

Mombaerts, P., Wang, F., Dulac, C., Chao, S.K., Nemes, A., Mendelsohn, M., Edmondson, J., and Axel, R. (1996). Visualizing an olfactory sensory map. Cell 87, 675–686.

Mosca, T.J., Hong, W., Dani, V.S., Favaloro, V., and Luo, L. (2012). Trans-synaptic Teneurin signalling in neuromuscular synapse organization and target choice. Nature 484, 237–241.

Paralkar, V.M., Vukicevic, S., and Reddi, A.H. (1991). Transforming growth factor beta type 1 binds to collagen IV of basement membrane matrix: implications for development. Dev Biol 143, 303–308.

Paralkar, V.M., Weeks, B.S., Yu, Y.M., Kleinman, H.K., and Reddi, A.H. (1992). Recombinant human bone morphogenetic protein 2B stimulates PC12 cell differentiation: potentiation and binding to type IV collagen. J Cell Biol 119, 1721–1728.

Piccioli, Z.D., and Littleton, J.T. (2014). Retrograde BMP signaling modulates rapid activity-dependent synaptic growth via presynaptic LIM kinase regulation of cofilin. J Neurosci 34, 4371–4381.

Prince, J.E., Brignall, A.C., Cutforth, T., Shen, K., and Cloutier, J.F. (2013). Kirrel3 is required for the coalescence of vomeronasal sensory neuron axons into glomeruli and for male-male aggression. Development 140, 2398–2408.

Prince, J.E., Cho, J.H., Dumontier, E., Andrews, W., Cutforth, T., Tessier-Lavigne, M., Parnavelas, J., and Cloutier, J.F. (2009). Robo-2 controls the segregation of a portion of basal vomeronasal sensory neuron axons to the posterior region of the accessory olfactory bulb. J Neurosci 29, 14211–14222.

Roos, J., Roos, M., Schaeffer, C., and Aron, C. (1988). Sexual differences in the development of accessory olfactory bulbs in the rat. J Comp Neurol 270, 121–131.

Schmidt, J.E., Suzuki, A., Ueno, N., and Kimelman, D. (1995). Localized BMP-4 mediates dorsal/ventral patterning in the early Xenopus embryo. Dev Biol 169, 37–50.

Shi, Y., and Massague, J. (2003). Mechanisms of TGF-beta signaling from cell membrane to the nucleus. Cell 113, 685–700.

Silvotti, L., Cavaliere, R.M., Belletti, S., and Tirindelli, R. (2018). In-vivo activation of vomeronasal neurons shows adaptive responses to pheromonal stimuli. Sci Rep 8, 8490.

Strittmatter, S.M., Valenzuela, D., Kennedy, T.E., Neer, E.J., and Fishman, M.C. (1990). G0 is a major growth cone protein subject to regulation by GAP-43. Nature 344, 836–841.

Strittmatter, S.M., Vartanian, T., and Fishman, M.C. (1992). GAP-43 as a plasticity protein in neuronal form and repair. J Neurobiol 23, 507–520.

Sudhof, T.C. (2017). Synaptic Neurexin Complexes: A Molecular Code for the Logic of Neural Circuits. Cell 171, 745–769.

Tadokoro, T., Gao, X., Hong, C.C., Hotten, D., and Hogan, B.L. (2016). BMP signaling and cellular dynamics during regeneration of airway epithelium from basal progenitors. Development 143, 764–773.

Taroc, E.Z.M., Prasad, A., Lin, J.M., and Forni, P.E. (2017). The terminal nerve plays a prominent role in GnRH-1 neuronal migration independent from proper olfactory and vomeronasal connections to the olfactory bulbs. Biol Open 6, 1552–1568.

Timmer, J.R., Wang, C., and Niswander, L. (2002). BMP signaling patterns the dorsal and intermediate neural tube via regulation of homeobox and helix-loop-helix transcription factors. Development 129, 2459–2472.

Wakabayashi, Y., and Ichikawa, M. (2007). Distribution of Notch1-expressing cells and proliferating cells in mouse vomeronasal organ. Neurosci Lett 411, 217–221.

Wang, X., Harris, R.E., Bayston, L.J., and Ashe, H.L. (2008). Type IV collagens regulate BMP signalling in Drosophila. Nature 455, 72–77.

Weiler, E., McCulloch, M.A., and Farbman, A.I. (1999). Proliferation in the vomeronasal organ of the rat during postnatal development. Eur J Neurosci 11, 700–711.

Yang, X., Li, C., Herrera, P.L., and Deng, C.X. (2002). Generation of Smad4/Dpc4 conditional knockout mice. Genesis 32, 80–81.

Love, M. I., Huber, W., and Anders, S. (2014) Moderated estimation of fold change and dispersion for RNA-seq data with DESeq2. Genome Biol. 15, 550

Zhou, Y., Zhou, B., Pache, L., Chang, M., Khodabakhshi, A. H., Tanaseichuk, O., Benner, C., and Chanda, S. K. (2019) Metascape provides a biologist-oriented resource for the analysis of systems-level datasets. Nature Communications. 10, 1523

